# Interspecies Interactions Drive Community-Level Selection in Microbial Coalescence

**DOI:** 10.64898/2026.02.11.704923

**Authors:** Jinyeop Song, Jiliang Hu, Jeff Gore

## Abstract

It has long been debated in ecology whether communities behave as cohesive units or as loose collections of independent species. Here, we study this question in the context of community coalescence, the mixing of previously isolated communities, using bacterial microcosm experiments combined with ecological modeling. Our results demonstrate that interspecies interaction strength determines whether communities or species are the units of selection during coalescence. When interactions are moderate to strong, one parental community consistently outcompetes the other, indicating community-level selection. In contrast, under weak interactions, species fates are uncorrelated and the two communities contribute equally to the coalesced outcome, indicating the absence of community-level selection. These patterns extend to communities derived from natural samples with greater taxonomic diversity and richness. Furthermore, we identify two distinct regimes underlying community-level selection in experiments with different media conditions: an emergent regime in which collective dynamics shape outcomes that cannot be predicted from species traits alone, and a top-down regime where dominant species determine the winning community. Together, these results reconcile conflicting observations on community-level selection during community coalescence by demonstrating that communities behave as cohesive units only when interactions are sufficiently strong.

## 1 Introduction

In nature, species coexist and interact within complex communities, yet whether these assemblages function as cohesive, integrated units or merely as loose collections of independently acting species remains a fundamental question in ecology. Historically, this tension has been framed through two influential paradigms. Clements’ “superorganism” view treats communities as discrete biological entities with emergent properties arising from species interdependencies, potentially as a result of coevolution ^1–5^. In contrast, Gleason’s individualistic hypothesis posits that species independently occupy niches, with community composition emerging from the coincidental overlap of species ranges ^6–9^. Despite decades of research, the conditions that determine when communities behave as cohesive units versus loose species assemblages remain unclear.

These contrasting frameworks make distinct predictions in the context of community coalescence, the mixing of previously isolated communities ^10,11^. Coalescence occurs across diverse contexts and scales: environmental disturbances trigger microbial community mixing in soils ^12,13^, flooding events promote coalescence in aquatic and estuarine habitats ^14^, skin microbiomes undergo exchange through daily social contact ^15^, and gut microbiomes are subject to wholesale community transfer through fecal microbiota transplantation ^16,17^. Coalescence brings species with distinct interaction histories into contact, creating novel cross-community interactions that can reshape the resulting assemblage ^10^. Thus, it provides a natural test for the individualistic versus holistic paradigms: The holistic view predicts that species within a community have correlated persistence, making the community the primary unit of selection, termed community-level selection. The individualistic view predicts species-level selection, yielding outcomes shaped by individual fitness regardless of community origin, with no systematic correlation in species persistence within parental communities.

Substantial theoretical and empirical evidence supports the holistic prediction that communities act as cohesive units during coalescence. Early theoretical work by Gilpin ^18^ suggested that pre-assembled communities, having already undergone internal competitive exclusion, possess structured interaction patterns that differ fundamentally from randomly assembled communities. Because species within such communities have been filtered to coexist, their survival outcomes become coupled, producing asymmetric post-coalescence communities dominated by one parental community ^19^. Notably, this structured interaction can arise due to ecological exclusion alone, even in the absence of long-term coevolution, as demonstrated in synthetic microbial communities ^20,21^. Building on this foundation, resource-consumer models formalized the mechanism by which such selection occurs, demonstrating that communities with superior collective fitness through resource consumption outcompete others in ways not predictable from individual species performance alone ^11,22–24^. Empirically, correlated selection between dominant and subdominant taxa has been observed across 100 coalescence experiments, confirming that species retention within communities is indeed correlated ^25^.

However, opposing evidence supports the individualistic prediction that species respond independently to coalescence events, with outcomes primarily determined by environmental sorting rather than collective community dynamics. Empirical work reports that coexistence between species from different source communities is common through niche partitioning rather than competitive exclusion during macro-scale biogeographic interchange in marine and terrestrial biomes ^26^; similar patterns have been observed in microbial systems where closely related species coexist via resource partitioning ^27^. Other work shows that species from the same community rarely go extinct together in the fossil record ^28^. In microbial systems, local environmental factors explain more variation than the presence of constantly interchanging neighboring communities ^29^. Strain-resolved metagenomic analyses of *in vitro* gut microbial communities showed species-level dynamics rather than community-level selection, with surviving species originating from both parental communities ^30,31^. These findings suggest that selection during coalescence acts on species-level traits rather than on emergent community properties.

Together, these contrasting lines of evidence indicate that communities behave as units of selection in some coalescence events but not others; however, the governing factors of these transitions remain poorly understood. Here, we combine empirical and theoretical approaches to investigate how interspecies interaction strength drives these contrasting outcomes. Using randomly assembled synthetic microbial communities across different interaction strengths, we classify post-coalescence outcomes into three types: Dominance (one community wins), Mixture (both persist), and Restructuring (a novel state emerges). Under weak interactions, Mixture prevails and species persistence is uncorrelated, consistent with individualistic dynamics. As interaction strength increases, the system shifts toward Dominance, where species persistence becomes correlated, indicating community-level selection. A theoretical model with minimal pairwise interactions reproduces these experimental observations, and the patterns generalize to natural communities with greater taxonomic complexity. Further analysis reveals two mechanistic regimes underlying community-level selection: one where a few dominant taxa largely determine the winner, and another where collective multi-species dynamics shape the outcome. These results reconcile conflicting observations by establishing interaction strength as the control parameter for community-level selection.

## 2 Results

### 2.1 Community-level selection is prevalent in microbial coalescence

Our central question is whether selection acts primarily on whole communities as cohesive units during community coalescence. To address this, we asked how often coalescence outcomes are represented primarily by one parental community versus blending of both or forming a novel community. Consider a coalescence event where communities A and B merge to produce community C (Fig. 1A). We represent each community by its normalized abundance vector and quantify the similarity between the coalesced community C and each parental community (Fig. 1B):

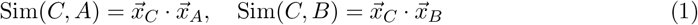

where 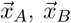 are normalized abundance vectors of parental communities A and B, and 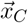 is that of the coalesced community. These two similarity scores yield a two-dimensional similarity map in which the location of C reflects how strongly it resembles each parental community. We partition this map into three coalescence outcome classes (see Methods): Dominance, where C closely resembles one parental community but not the other; Mixture, where C similarly resembles both parental communities; and Restructuring, where C diverges from both parental communities into a novel configuration. Crucially, Dominance reflects community-level selection: one parental community displaces the other while retaining its internal compositional structure, indicating that its species persist collectively.

**Figure 1:**
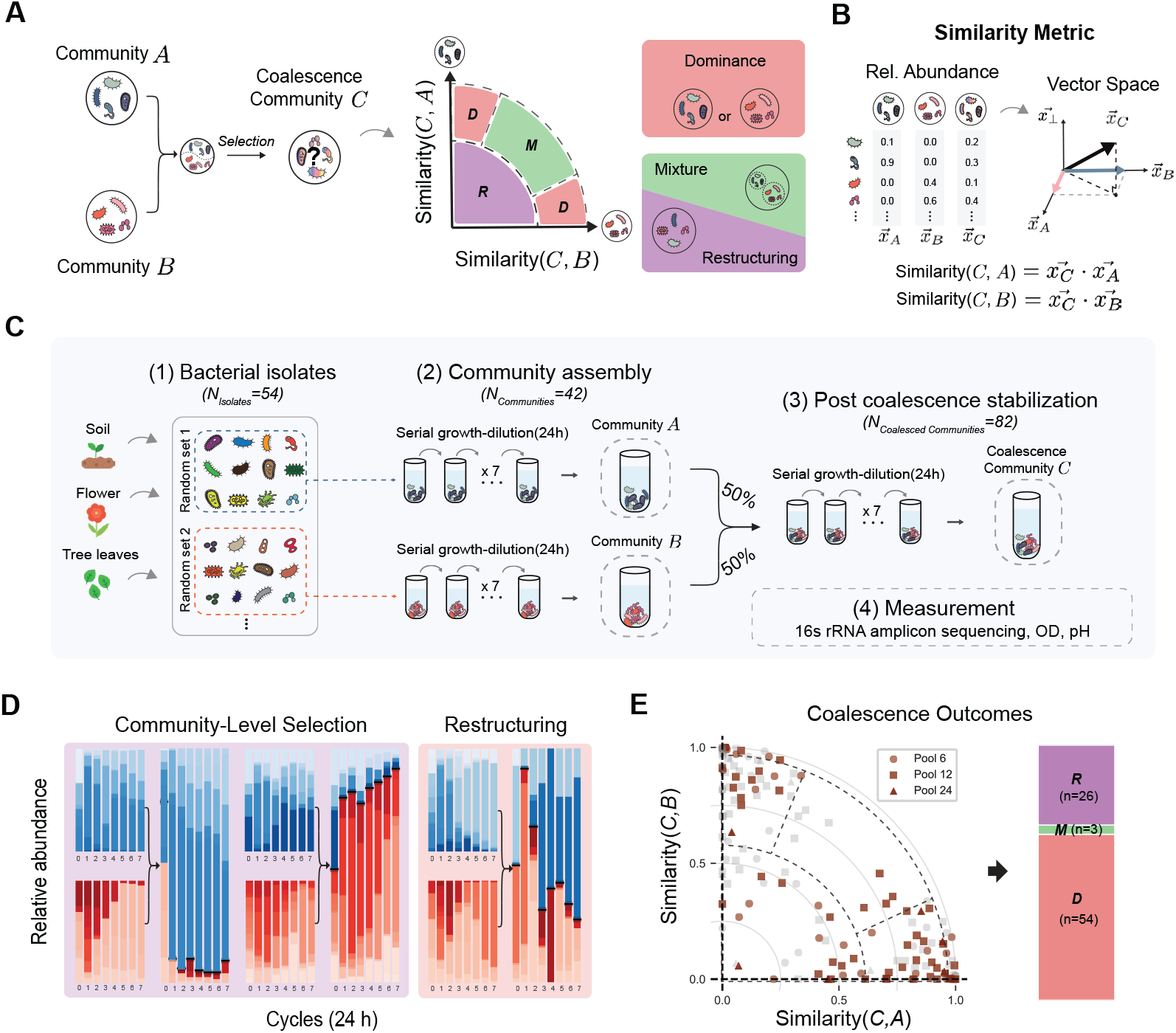
Fig. 1. Coalescence of synthetic microbial communities frequently yields Dominance. **a**, Schematic of coalescence: parental communities A and B are mixed to produce C. Outcomes are classified as Dominance (one parent wins), Mixture (both persist), or Restructuring (novel state). **b**, We use similarity to quantify compositional resemblance between communities and define a two-dimensional similarity map for classifying outcomes. **c**, Experimental workflow: 30 parental communities (initial richness 6, 12, or 24) were assembled from 54 bacterial isolates, stabilized, mixed pairwise (n = 83), and restabilized. **d**, Representative time courses showing Dominance (left, center) and Restructuring (right). **e**, Dominance is the most frequent outcome in Base medium. Of 83 coalescence events, 54 (65%) were classified as Dominance, 26 (31%) as Restructuring, and 3 (4%) as Mixture. Different symbols indicate initial richness (circles: 6; squares: 12; triangles: 24 species).

To quantify the occurrence of outcome types experimentally, we assembled random in-vitro communities and subjected them to pairwise coalescence (Fig. 1C). We first curated a strain library of 54 bacterial isolates collected from diverse environments (soil, tree surface, and flower stamen). The library is phylogenetically broad, spanning 29 families across three phyla: Proteobacteria, Firmicutes, and Bacteroidota (Extended Data Fig. 1; Supplementary Fig. 1). From this library, we assembled 30 parental communities with varying initial richness (6, 12, or 24 species; see Methods) and stabilized them for 7 days under daily growth–dilution cycles (×30 every 24 h) (Fig. 1C). Our initial experiments were done in Base medium (1 g L^−1^ yeast extract, 1 g L^−1^ soytone, 10 mM sodium phosphate, 0.1 mM CaCl_2_, 2 mM MgCl_2_, 4 mg L^−1^ NiSO_4_, 50 mg L^−1^ MnCl_2_, 5 g L^−1^ glucose and 4 g L^−1^ urea; pH 6.5), a buffered complex medium that we have previously determined has moderate interaction strength and supports high species coexistence (Supplementary Information, mean species survival ratio = 74 ± 2%). We then performed 83 pairwise coalescence events by mixing stabilized parental communities at equal volume and restabilizing for an additional 7 days. Community compositions of parental and coalesced communities were measured at the end of stabilization by 16S rRNA amplicon sequencing (Amplicon Sequence Variants, ASVs), and we also recorded the optical density (OD_600_) and pH of the communities to contextualize assembly and post-coalescence dynamics.

Representative time series illustrate the spectrum of post-coalescence dynamics (Fig. 1D). In outcomes classified as Dominance, one parental lineage rapidly displaces the other after mixing and the coalesced community converges to a composition closely matching that parental community, while largely preserving its internal relative-abundance structure over serial dilution cycles. In the Restructuring case, the merged community converges toward a novel state distinct from both parental communities. Among 3 representative time-series trajectories, we did not observe Mixture cases in which both parental communities remained comparably represented after coalescence (see Supplementary Fig. 26).

Projecting all outcomes into the similarity space based on stabilized compositions (Fig. 1E) revealed that Dominance is the most frequent empirical outcome. Of the 83 coalescence events, 54 were classified as Dominance, 26 as Restructuring, and 3 as Mixture (Fig. 1E). This pattern of Dominance as the most frequent outcome was robust across variants of similarity metrics (Extended Data Fig. 2). One possibility is that the observed frequency of Dominance results from skewed species abundance distributions rather than correlated selection among species from the same parental community. To rule out this possibility, we compared experimental outcomes against two null models that assume no correlation in species selection: (1) abundance-weighted random selection and (2) shuffled abundance (see Supplementary Information). The experimentally observed asymmetry significantly exceeded both null expectations (Extended Data Fig. 3), supporting that Dominance reflects correlated species selection within parental communities.

### 2.2 Theoretical model with random interactions reproduces community-level selection

To gain insight into why Dominance is the prevalent outcome, we introduced a generalized Lotka– Volterra (gLV) model ^32,33^ that mirrors the experimental protocol (Fig. 2A). In this classic model, species grow logistically and compete pairwise:

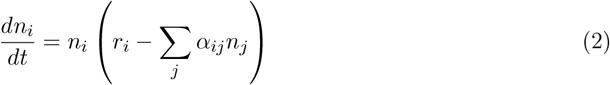

where n_*i*_ is abundance, r_*i*_ is growth rate, and α_*ij*_ is the competition coefficient between species i and j (with self-interaction α_*ii*_ = 1). We fixed r_*i*_ = 1 and drew off-diagonal competition coefficients from a uniform distribution 𝕌(0, 2*µ*) with mean *µ*, which controls the average competition strength: higher *µ* means stronger interspecies competition ^34,35^. From a pool of 54 species, we generated two parental communities of 12 species each with no shared species, allowed each to reach equilibrium, then mixed them equally to simulate coalescence (Fig. 2A; see Methods for details). Post-coalescence compositions were analyzed using the same similarity metrics as in the experiments.

**Figure 2:**
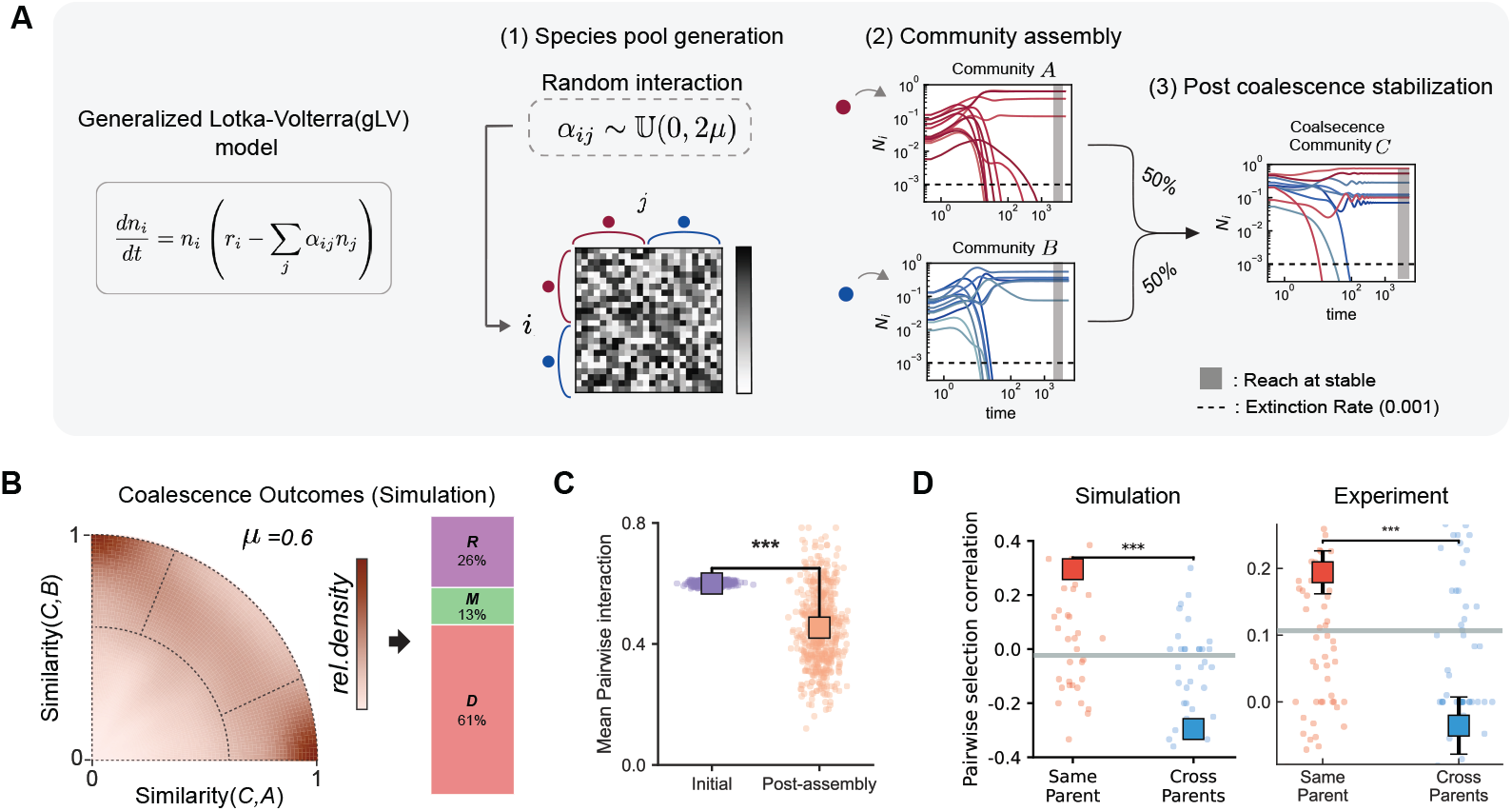
Fig. 2. Generalized Lotka–Volterra model of community coalescence. **a**, Simulation workflow: species grow according to the generalized Lotka–Volterra (gLV) equations, where interaction coefficients α_*ij*_ are drawn from a uniform distribution 𝕌(0, 2*µ*) with mean *µ* controlling mean interaction strength. Two equilibrated parental communities are mixed and simulated to steady state. **b**, Outcome density map (*µ* = 0.6, n = 1,200 simulations). The model reproduces the high frequency of Dominance (61%) observed experimentally. **c**, Community assembly significantly reduces the mean interaction strength among surviving species (paired t-test, p < 0.001). **d**, Pairwise selection correlation. Species pairs from the same parental community show positive correlation (co-survival), while cross-community pairs show negative correlation, indicating community-level selection. This pattern holds for both simulations (left) and experiments (right). Error bars, s.e.m.

We simulated 1,200 random coalescence events at interaction strength *µ* = 0.6 and analyzed the similarity of each coalescence outcome to the two parental communities. At this interaction strength, the random-interaction model quantitatively reproduces the high frequency of Dominance observed in the experiments. Outcomes concentrate near the axes rather than the diagonal, indicating asymmetric dominance by one parental community (Fig. 2B). Quantitatively, Dominance accounts for 61% of all outcomes, far more frequent than Restructuring (26%) or Mixture (13%; Fig. 2B, right). This observation suggests that interspecies interactions alone, a minimal and generic condition, are sufficient to reproduce the high frequency of Dominance observed in coalescence.

To understand how Dominance emerges in the model and whether it reflects community-level selection, we examined the role of the assembly process. During assembly, competitive exclusion filters out species with strong competitive interactions, leaving communities with reduced mean interaction strength compared to the initial pool (paired t-test, P < 0.001; Fig. 2C). Because surviving species compete weakly with each other but face stronger competition from outsiders, their fates become coupled during coalescence: they tend to persist or go extinct together. We quantified this effect using pairwise selection correlation—the degree to which species pairs share the same survival outcome (both persist or both go extinct) across coalescence events (Supplementary Information; Fig. 2D). Species from the same parental community showed strongly positive correlations, while cross-community pairs showed negative correlations—a pattern observed in both simulations and experiments (Fig. 2D). This confirms that member species within a parental community, having undergone assembly together, have coupled fates and are coselected during coalescence. Together, the prevalence of Dominance combined with positive within-community selection correlation provides evidence for community-level selection in this system.

### 2.3 Interaction strength controls coalescence outcome type and degree of community-level selection

Having established that random interspecies interactions can recapitulate Dominance with correlated pairwise selection, we next investigated how interaction strength alters coalescence outcomes. We simulated 1,200 coalescence events at each of three representative interaction strengths (*µ* = 0.3, 0.6, 0.8) and mapped outcomes into the similarity space (Fig. 3A). At weak interactions (*µ* = 0.3), outcomes clustered in the interior of the map, indicating Mixture; as *µ* increased, outcomes shifted toward the axes, indicating Dominance. We quantified this shift using the Parental Dominance Index (PDI), which captures the relative contribution of each parental community (0: parental community B, 0.5: equal, 1: parental community A; see Methods). PDI distributions shifted from unimodal near 0.5 at *µ* = 0.3 to bimodal at higher *µ* (Fig. 3A), reflecting a transition in which Dominance becomes increasingly frequent, indicating community-level selection.

**Figure 3:**
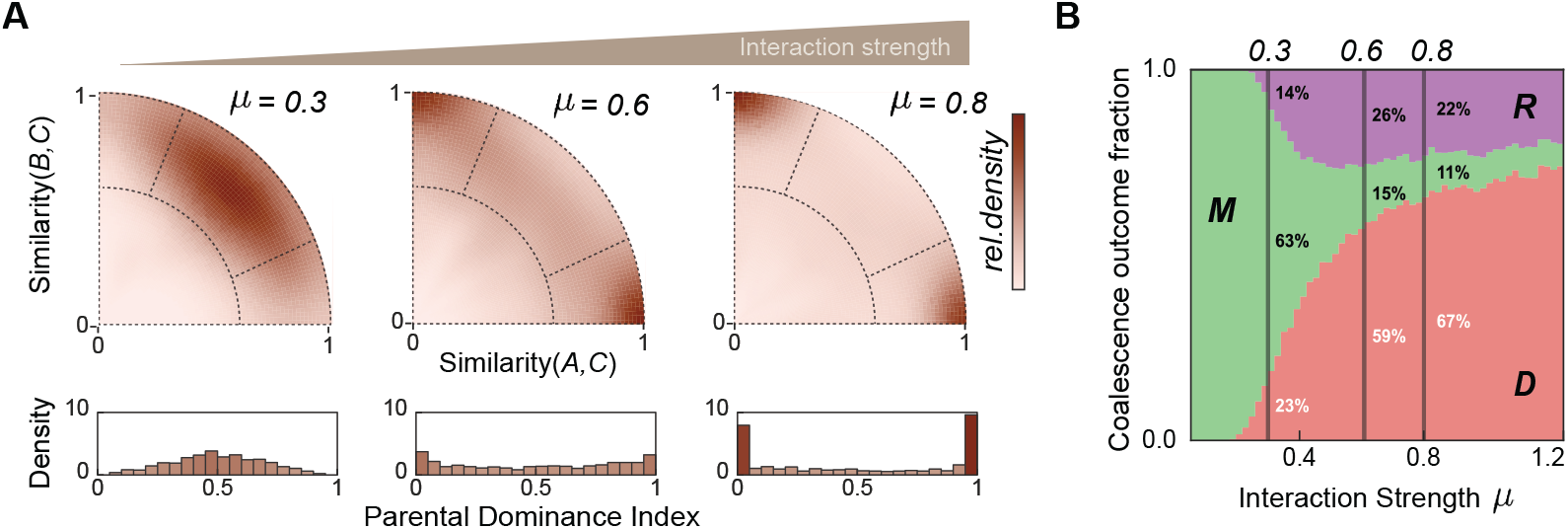
Fig. 3. Interaction strength controls the transition between coalescence outcome types in simulation. **a**, Simulated coalescence outcome maps at three representative interaction strengths: *µ* = 0.3, 0.6, 0.8; n = 1,200 simulations per condition. Outcomes shift from central Mixtures at weak interactions (*µ* = 0.3) to axis-aligned Dominance at strong interactions (*µ* = 0.8). Histograms show the Parental Dominance Index (PDI) transitioning from unimodal to bimodal. **b**, Outcome fractions across interaction strengths (*µ* = 0–1.2). Mixture predominates at low *µ*, while Dominance becomes prevalent as *µ* increases. Restructuring peaks at intermediate strengths. Error bars, s.e.m.

Across interaction strengths from *µ* = 0 to 1.2, we observed a systematic shift in coalescence outcomes (Fig. 3B), with Mixture predominating at weak interactions and Dominance at strong interactions. Restructuring emerged at moderate to high interaction strengths. These patterns were robust to variation in carrying capacities, interaction distributions, similarity metrics, and community size (Supplementary Figs. 2–4; Extended Data Fig. 4). Notably, the frequency of Dominance remained relatively stable across parental communities ranging from 4 to 48 species, spanning our experimental species pool range. To determine whether this transition from Mixture to Dominance corresponds to the degree of co-selection, we examined pairwise selection correlation across interaction strengths. Notably, within-community pairwise selection correlation emerged only at high interaction strengths, indicating that species fates become coupled only when interactions are sufficiently strong (Extended Data Fig. 5). These findings indicate that interaction strength simultaneously determines both coalescence outcome type and the level at which selection operates.

### 2.4 Nutrient-dependent interaction strength in experiments recapitulates model predictions

Following prior work showing that nutrient concentration intensifies microbial competition ^34–36^, we conducted additional coalescence experiments by removing or augmenting glucose and urea in the Base medium used in Fig. 1 (Methods). This yielded two additional media conditions (Fig. 4A): Nutr− (no added glucose/urea) and Nutr+ (high supplementation). Higher nutrient concentration amplifies consumer–resource feedbacks and intensifies environmental modification, thereby strengthening interspecies interactions ^36^. To empirically validate this effect, we measured failed invasion frequency using pairwise invasion assays among the 12 most abundant isolates (95:5 initial frequency; Methods, Supplementary Figs. 5–7). The fraction of failed invasions increased monotonically with nutrient supply (Nutr−: 2 ± 1%; Base: 33 ± 4%; Nutr+: 48 ± 4%; mean ± s.e.m.; Fig. 4B). Assuming the uniform distribution used in the model, calibrating these values against gLV simulations yielded approximate mean interaction strengths of *µ* ≈ 0.5 for Nutr−, *µ* ≈ 0.7 for Base, and *µ* ≈ 0.9 for Nutr+ (see Supplementary Methods).

**Figure 4:**
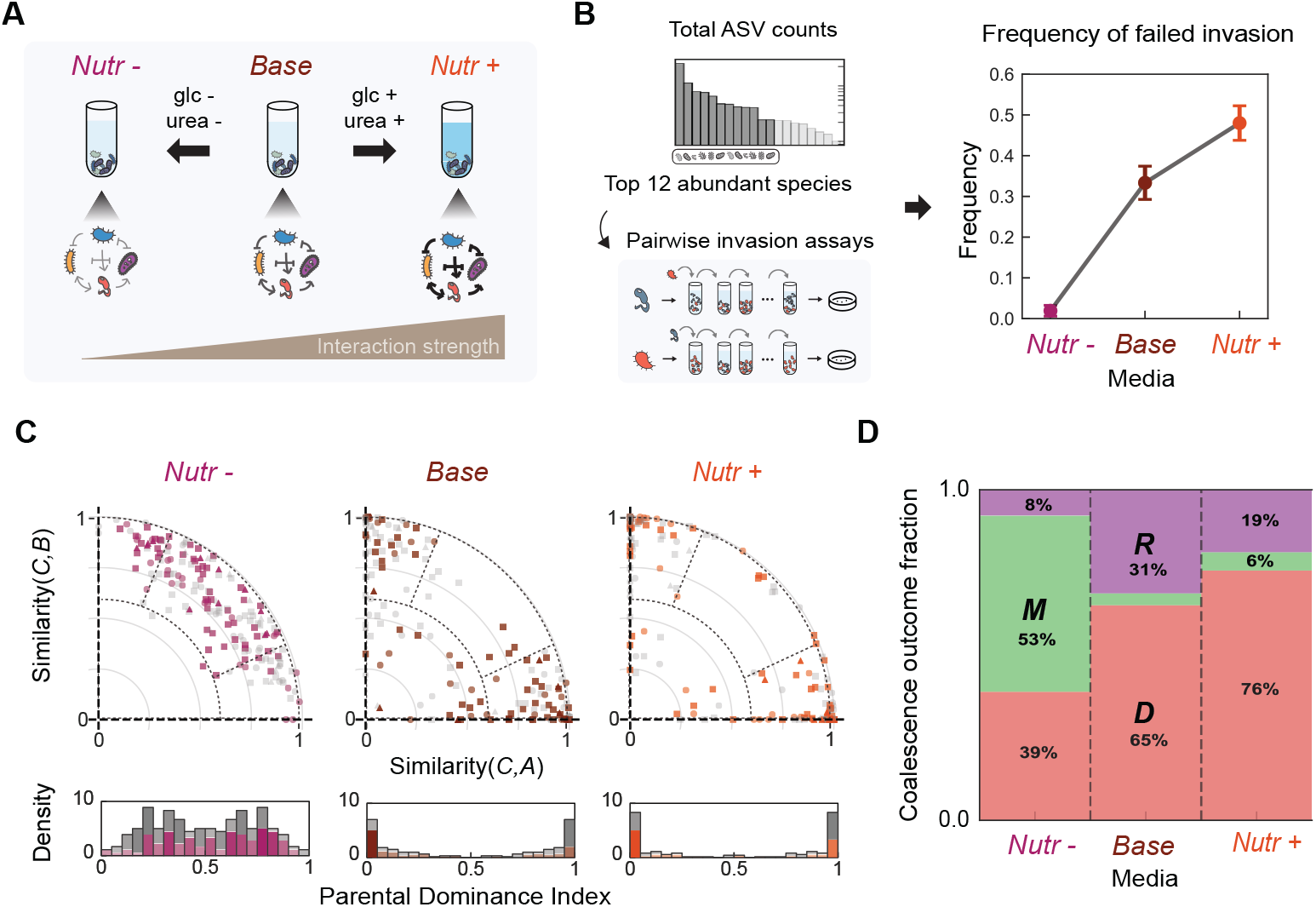
Fig. 4. Nutrient concentration modulates interaction strength and validates model predictions. Experimental manipulation of nutrient concentration confirms that stronger interactions shift coalescence outcomes from Mixture toward Dominance. **a**, Schematic of nutrient manipulation: we varied glucose and urea concentrations to create three media conditions with increasing interaction strength (Nutr−, Base, Nutr+). **b**, Pairwise invasion assays confirm that nutrient concentration modulates interaction strength. Failed invasion frequency among the 12 most abundant isolates (95:5 initial frequency, n = 132 assays per medium) increases monotonically with nutrient concentration (Nutr−: 2 ± 1%, Base: 33 ± 4%, Nutr+: 48 ± 4%; mean ± s.e.m.). **c**, Coalescence outcome distributions shift systematically with nutrient concentration. Scatter plots show outcomes in similarity space for each medium (Nutr−: n = 90, magenta; Base: n = 83, brown; Nutr+: n = 90, orange); different symbols indicate initial richness. Histograms show PDI distributions. **d**, The fraction of Dominance increases with nutrient concentration (39% in Nutr−, 65% in Base, 76% in Nutr+) while Mixture declines from 53% to 4% to 6% (chi-square test for trend, p < 0.001), consistent with model predictions. Error bars, s.e.m.

Given that nutrient concentration modulates interaction strength, we performed coalescence experiments in Nutr− and Nutr+ media using the same parental community library to examine how interaction strength affects coalescence outcomes. The distribution of outcomes in similarity space shifted systematically with nutrient concentration (Fig. 4C), which we quantified by the fraction of each outcome type (Fig. 4D). In Nutr− (n = 90), where interactions are weakest, Mixture was the most frequent outcome (53%), and Dominance was substantially reduced to 39% compared to the Base medium (Dominance: 65%, Mixture: 4%). In Nutr+ (n = 90), Dominance further increased to 76%. Pairwise selection correlation analysis confirmed that this shift corresponds to a transition in the level of selection (Extended Data Fig. 6). In Base and Nutr+ media, within-community species pairs showed significantly higher selection correlation than cross-community pairs (P < 0.001), consistent with community-level selection. In Nutr−, no such correlation was observed, indicating species-level dynamics. These results experimentally validate the theoretical prediction: weaker interactions yield Mixture with uncorrelated species fates, while stronger interactions yield Dominance with community-level selection.

### 2.5 Dominant community predictability reveals two mechanistic regimes

We observed Dominance in both Base and Nutr+ media, consistent with community-level selection as confirmed by pairwise selection correlation (Extended Data Fig. 6). We next asked whether we can predict which parental community will win during coalescence. Prior work has suggested that species-level competitive outcomes may correlate with Dominance direction ^10,37^. Inspired by these observations, we tested whether pairwise competition between dominant species ^34,36^ predicts which community wins (Fig. 5A).

**Figure 5:**
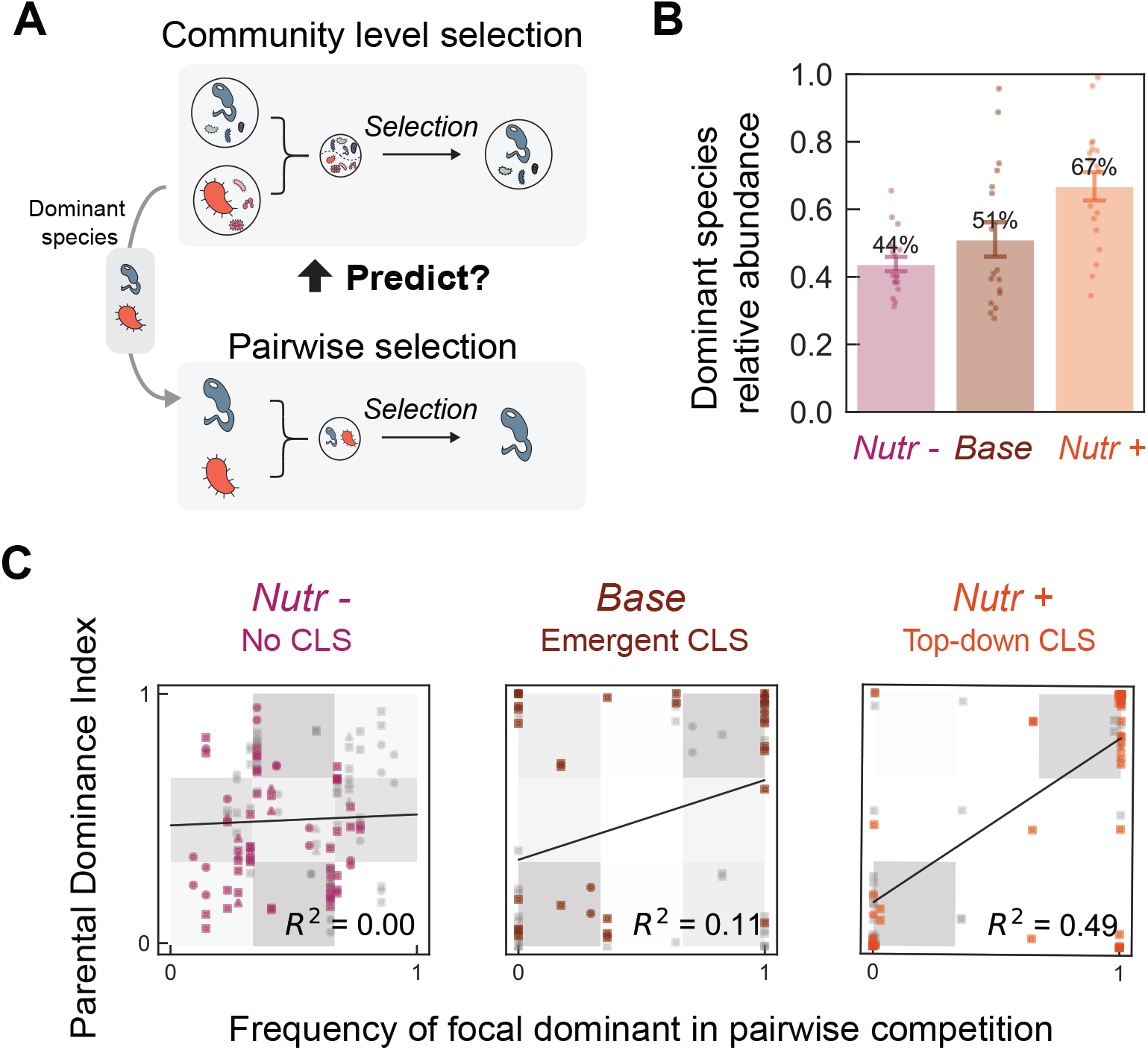
Fig. 5. Predictability of Dominance direction reveals two mechanistic regimes. The direction of Dominance (which parental community wins) becomes increasingly predictable from dominant species competition across media conditions, distinguishing an emergent regime (Base) from a top-down regime (Nutr+). **a**, Schematic: can pairwise competition between dominant species predict which community wins? **b**, Relative abundance of dominant species increases with nutrient concentration (Nutr−: 44 ± 2%; Base: 51 ± 5%; Nutr+: 67 ± 4%; mean ± s.e.m.). **c**, Dominant species competitive success versus PDI of coalescence outcomes; gray shading indicates Dominance. Predictive power: R^2^ = 0.00 in Nutr−, R^2^ = 0.11 (p = 0.03) in Base (emergent regime), and R^2^ = 0.49 (p < 0.001) in Nutr+ (top-down regime). Linear regression with 95% confidence intervals shown.

The relative abundance of dominant species in stabilized parental communities increased with nutrient concentration (44 ± 2% in Nutr−, 51 ± 5% in Base, 67 ± 4% in Nutr+; mean ± s.e.m., Fig. 5B), suggesting that dominant species exert greater influence under higher nutrient concentration. We therefore tested whether pairwise competition between dominant species predicts the coalescence outcome by analyzing the linear relationship between dominant-species competitive success (from invasion assays) and the PDI of coalescence outcomes (Fig. 5C). Predictive power varied markedly across media: in Nutr−, where Mixture predominates and pairwise selection correlation is weak, pairwise competition showed no predictive power (R^2^ = 0.00); in Base medium, predictive power was weak (R^2^ = 0.11), consistent with collective multi-species dynamics shaping outcomes; in Nutr+, pairwise competition was more predictive (R^2^ = 0.49), indicating that the dominant species contributes substantially to determining which community wins. These results suggest that community-level selection spans a mechanistic continuum: at one end, an emergent regime (Base medium) where multi-species dynamics collectively shape the outcome and no single species trait is predictive, and at the other, a top-down regime (Nutr+ medium) where a few dominant taxa determine the winner.

We explored the mechanistic basis of the top-down regime by examining environmental modification through pH. In our system, dominant species are often strong pH modifiers that either acidify or alkalinize the medium (Supplementary Fig. 8). In both Base and Nutr+ media, when these pH-modifying species become dominant within a community, they determine the community’s overall pH; communities dominated by acidifiers become acidic, while those dominated by alkalizers become alkaline (Supplementary Fig. 9). In coalescence events between acidic (pH < 6.5) and alkaline (pH > 7.5) parental communities, the acidic community won in only 56% of cases in Base medium (n = 41), but won in 91% of cases in Nutr+ medium (n = 32, Fisher’s exact test p < 0.0001; Extended Data Fig. 8), consistent with the hypothesis that high nutrients amplify metabolic activity, thereby intensifying pH modification and excluding pH-sensitive species ^36,38^. Thus, in Nutr+ medium, the dominant species determines community pH, and community pH predicts coalescence outcome, providing a mechanistic explanation for the top-down regime.

### 2.6 Interaction-dependent coalescence outcomes generalize to natural communities

The synthetic communities described above were assembled from individually isolated bacterial strains, allowing precise control over initial composition but raising the question of whether these patterns generalize to more ecologically realistic settings. To address this, we performed coalescence experiments using communities derived from natural environmental samples, which harbor complex species assemblages shaped by evolutionary and ecological processes in their natural habitats.

We collected six environmental samples from diverse microhabitats (soil, compost, and decomposing organic matter) and established bacterial communities through seven rounds of serial growth-dilution in laboratory media across all three nutrient conditions (Nutr−, Base, and Nutr+; Fig. 6A). After stabilization, these natural sample-derived communities exhibited higher ASV richness than synthetic communities (mean of 13.7 ± 7.2 ASVs above 0.1% threshold, compared to 9.8 ± 4.8 in synthetic communities) and low ASV overlap among communities from different samples (Supplementary Figs. 22–25).

**Figure 6:**
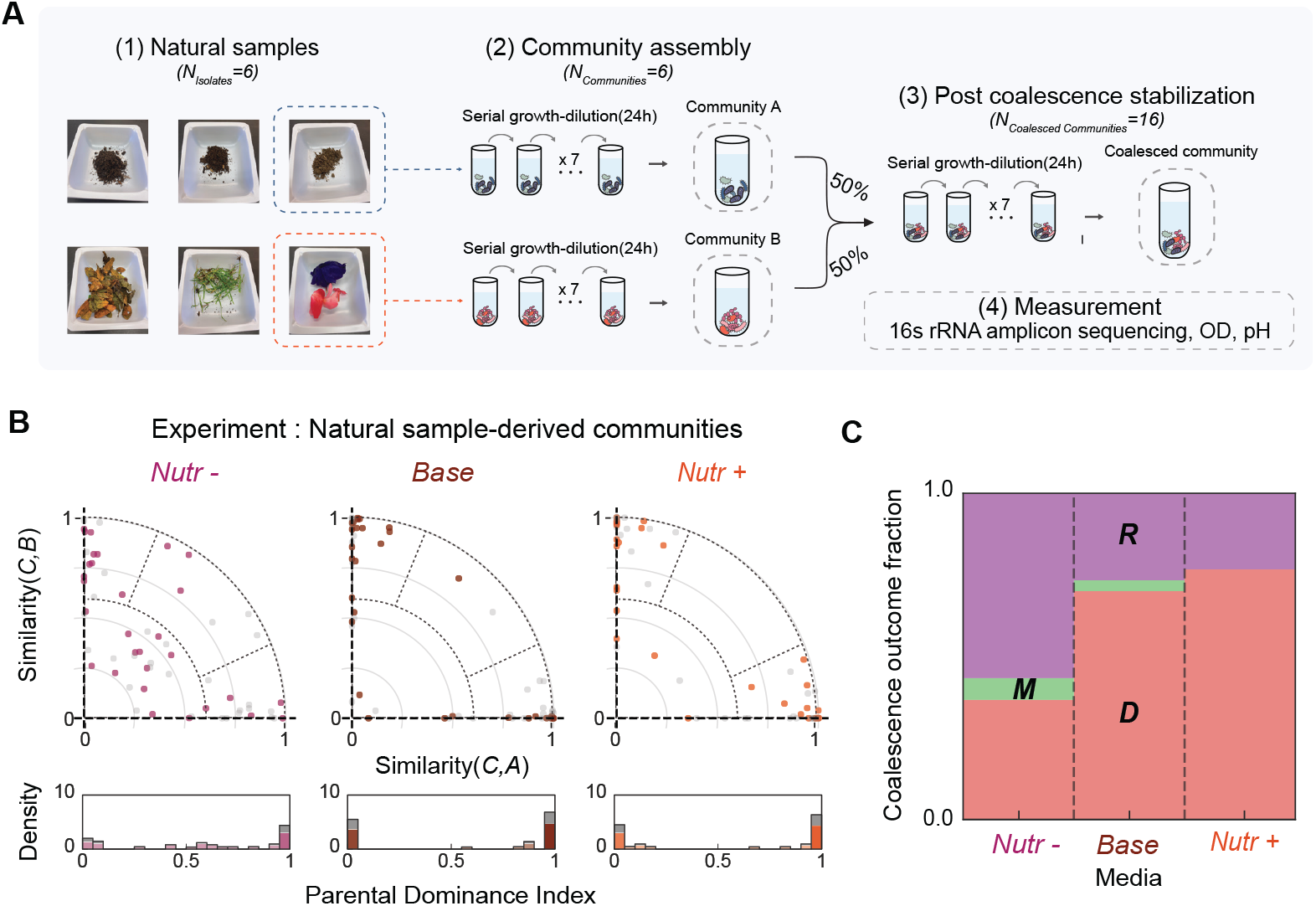
Fig. 6. Dominance generalizes to natural sample-derived communities. Natural environmental communities show qualitatively similar coalescence patterns to synthetic communities, with Dominance frequency increasing under higher nutrient concentration. **a**, Experimental workflow follows Fig. 1c, but uses natural sample-derived communities from six environmental samples including soil, compost, and decomposing organic matter. **b**, Coalescence outcome distributions in similarity space for natural communities across three media conditions (n = 30 coalescence events per condition, 15 unique pairs × 2 replicates). Similar to synthetic communities, outcomes cluster near the diagonal under weak interactions and shift toward the axes under strong interactions. Histograms below show PDI distributions. **c**, The fraction of Dominance outcomes increases with nutrient concentration in natural communities (37% in Nutr−, 70% in Base, 77% in Nutr+), consistent with patterns observed in synthetic communities. Error bars, s.e.m.

We performed 15 pairwise coalescence events with two biological replicates each (n = 30 per condition) across all three nutrient conditions after stabilization and analyzed outcomes using the same similarity framework applied to synthetic communities. Natural sample-derived communities showed similar patterns to synthetic communities: outcomes clustered along the axes in similarity space (Fig. 6B,C). The fraction of Dominance outcomes increased with nutrient concentration (37% in Nutr−, 70% in Base, 77% in Nutr+), mirroring the interaction-strength dependence observed in synthetic communities. Natural communities showed higher Restructuring fractions, possibly reflecting greater taxonomic diversity and more complex interaction networks. These results demonstrate that the interaction-dependent shift toward community-level selection is robust and generalizes beyond precisely controlled synthetic consortia to naturallyevolved microbial assemblages.

## 3 Discussion

Our work demonstrates that interspecies interaction strength governs community coalescence outcomes, determining both the dominant outcome type and the level at which selection operates. In both experiments and modeling, when interactions are strong, Dominance prevails and species persistence is correlated within parental communities, indicating community-level selection; when interactions are weak, Mixture predominates and species respond independently. Furthermore, predictability from dominant species reveals two distinct mechanisms underlying Dominance: a top-down regime driven by dominant species via pH modification, and an emergent regime where collective dynamics reduce predictability. These findings reconcile the Clements–Gleason debate: interspecies interaction strength determines whether communities behave as cohesive units or loose species assemblages.

Our work provides the first experimental demonstration of alternative regimes in community coalescence, reconciling previously contrasting observations of community-level versus specieslevel selection. Prior studies have reported both outcomes: some found communities behaving as cohesive units ^22,25^, while others observed species responding independently ^26,29^. Our results suggest that these conflicting observations may reflect differences in interaction strength across experimental systems. For instance, the absence of community-level selection in *in vitro* gut microbiome coalescence experiments ^30,31^ is consistent with our framework. Rich media such as BHI are analogous to our Nutr− condition, providing complex and diverse resources while lacking high concentrations of readily metabolizable carbon sources. This resource structure reduces both direct resource competition and pH-mediated interactions, thereby weakening interspecies interactions ^36^ and favoring species-level rather than community-level dynamics. Likewise, interaction strength may vary systematically across ecosystems: microbial biofilms experience strong metabolic interactions due to high cell densities and shared resources ^39^, whereas mobile macroscopic organisms interact more transiently across larger spatial scales. These differences may explain why community-level selection is more frequently observed in microbial coalescence studies. This framework provides a unified perspective: rather than asking whether communities are cohesive units, we should ask under what conditions they become so, with interaction strength emerging as a key determinant.

A growing body of work has identified interaction strength as a key control parameter governing diverse aspects of microbial community dynamics, including diversity ^34,40^, stability ^36,41^, coexistence ^33^, and invasion outcomes ^35,42^. Our results extend this claim to include coalescence, providing further evidence that interaction strength determines when communities exhibit collective behavior. The parallel with priority effects is particularly instructive; recent work showed that assembly order shapes final composition only under strong interactions, whereas weak interactions lead to convergent outcomes regardless of assembly history ^35,43^. Our coalescence results mirror this pattern—strong interactions preserve parental identity, while weak interactions yield convergent Mixtures. Together, these findings reinforce interaction strength as a coarse-grained parameter that predicts history-dependent community dynamics across multiple ecological contexts.

In nature, coalescence occurs in richer settings that may shift regimes and outcomes. Environmental heterogeneity (temperature, pH buffering, and other physicochemical factors) covaries with interaction strength and can reshape coalescence across habitats ^10,37,44,45^. In hostassociated microbiomes, host filtering, priority effects, and spatial structure further influence successful colonization, making coalescence a useful framework for predicting and steering microbiome transfer ^14,46,47^. Our experiments focus on steady-state outcomes and thus do not capture temporal dynamics such as fluctuating resources, migration pulses, or host responses. Additionally, both our experimental system and theoretical model are dominated by competitive interactions; systems with substantial mutualism or facilitation may exhibit qualitatively different dynamics. Incorporating these axes into both theory and experiment will help build a more comprehensive picture of community-level selection across habitats and scales.

## 4 Methods

### Microbial Strain Library and Culture Conditions

We used a library of 54 bacterial isolates from environmental samples (soil, tree surface, and flower stamen) collected in Cambridge, MA, USA, spanning 29 families across three phyla (Proteobacteria, Firmicutes, and Bacteroidota; Extended Data Fig. 1; Supplementary Fig. 1). Isolates were purified by serial streaking and stored as glycerol stocks at −80 °C. Experiments were conducted in culture medium containing 1 g L^−1^ yeast extract, 1 g L^−1^ soytone, 10 mM sodium phosphate, 0.1 mM CaCl_2_, 2 mM MgCl_2_, 4 mg L^−1^ NiSO_4_, 50 mg L^−1^ MnCl_2_ (pH 6.5), supplemented at three nutrient levels—Nutr− (no added glucose/urea), Base (5 g L^−1^ glucose, 4 g L^−1^ urea), and Nutr+ (20 g L^−1^ glucose, 16 g L^−1^ urea)—which produce different strengths of interspecific interactions. Communities were grown in 300 *µ*L volumes in 96-well deep-well plates at 25 °C with shaking at 800 rpm. Full media composition and culture conditions are provided in Supplementary Methods.

### Community Assembly and Coalescence Experiments

Parental communities (n = 30) were assembled at three richness levels (6, 12, or 24 species) by sequential assignment of isolates from the strain library, each with two biological replicates. For 6-species and 12-species parental communities, non-overlapping sets were assembled (e.g., community 1: strains 1–6, community 2: strains 7–12). Communities were stabilized through seven daily serial dilutions (×30) before coalescence. Coalescence experiments mixed two pre-stabilized parental communities at 1:1 volume ratio, followed by seven additional serial transfers to reach new steady states (Fig. 1C). In total, 83 pairwise coalescence events were performed in Base medium, and additional experiments were conducted across all three nutrient conditions. Full experimental details are provided in Supplementary Methods.

### 16S rRNA Sequencing

Community composition was measured by 16S rRNA amplicon sequencing (V4 region). DNA was extracted using QIAGEN DNeasy PowerSoil kit, and sequencing was performed at Argonne National Laboratory. Amplicon sequence variants (ASVs) were identified using DADA2^48^ with SILVA 138^49^ as reference. Species richness was defined as ASVs with ≥0.1% relative abundance, corresponding to the extinction threshold used in simulations. Raw sequencing reads are available at Dryad (http://datadryad.org/share/LINK_NOT_FOR_PUBLICATION/kQACU7LCmQclVZfGZk0bS5ZPUVL_grhwah2zvFY4m9s). Full sequencing and data processing details are provided in Supplementary Methods.

### Optical Density and pH Measurements

Optical density (OD_600_) was measured after each 24-hour growth cycle using a plate reader (BioTek Synergy H1). Community pH was measured using a benchtop pH meter (Apera Instruments PH5500). These measurements were used to monitor community growth dynamics and to assess environmental modification by dominant species. Full measurement protocols are provided in Supplementary Methods.

### Classification of Coalescence Outcomes

Coalescence outcomes were classified based on similarity between the coalesced community and each parental community. Each community was represented as a normalized abundance vector (see Supplementary Methods for normalization details), and similarity was computed as the dot product between the coalesced community vector and each parental vector. These two similarity scores place each outcome in a two-dimensional similarity space. From the similarity scores, we derived: (1) retention magnitude, quantifying how much of the coalesced composition is explained by the parental communities; and (2) parental dominance index (PDI), quantifying selection preference toward one parental community (0: parental community B dominance, 0.5: equal contributions, 1: parental community A dominance). Outcomes were categorized as Restructuring (low retention; substantial ecological reorganization), Mixture (high retention with PDI near 0.5; balanced parental contributions), or Dominance (high retention with PDI near 0 or 1; one parental community overwhelms the other). To assess whether Dominance reflects community-level selection, we computed pairwise selection correlations measuring whether species from the same parental community share selection outcomes during coalescence. Full mathematical framework is provided in Supplementary Methods.

### Lotka–Volterra Simulations

Community dynamics were modeled using generalized Lotka–Volterra (gLV) equations (Eq. 2). We fixed growth rates to 1 and self-interaction coefficients α_*ii*_ = 1, and drew off-diagonal competition coefficients from a uniform distribution U(0, 2*µ*) with mean *µ*, which controls mean interaction strength. From a pool of 54 species, we randomly generated two parental communities of 12 species each with no shared species. Communities were equilibrated numerically (species below 0.1% relative abundance threshold set to zero), then mixed pairwise at equal proportions to simulate coalescence. Each simulation used independently sampled interaction matrices to explore diverse ecological contexts. Full simulation parameters and robustness analyses are provided in Supplementary Methods.

### Pairwise Invasion Assays

To empirically estimate interaction strength, we performed pairwise invasion assays among the 12 most abundant isolates. Each pair was tested in both directions (resident:invader = 95:5) and propagated through seven daily dilution cycles across all three nutrient conditions. Final compositions were determined by colony counting. An invasion was scored as failed if the invader remained below 1% relative abundance. Pairwise outcomes were then classified as coexistence (both isolates above 10% in both directions), exclusion (the same isolate drove its competitor below 1% in both directions), or bistability (each isolate excluded the other when resident). The fraction of failed invasions served as a proxy for mean interaction strength (Fig. 4B). Full experimental details are provided in Supplementary Methods.

### Natural Sample-Derived Communities

We performed coalescence experiments using communities derived from six natural environmental samples (soil, compost, decomposing organic matter) collected in Cambridge, MA. Unlike synthetic communities assembled from isolated strains, these communities retain complex species assemblages from their native habitats. Samples were enriched through seven serial dilution cycles to establish stable communities, then 15 pairwise coalescence events (each with two biological replicates, n = 30 per condition) were conducted across all three nutrient conditions. Full experimental details are provided in Supplementary Methods.

### Statistical Analyses

Statistical significance was defined at p < 0.05. Paired t-tests compared mean interaction strengths before and after assembly. Permutation tests (1,000 permutations) assessed pairwise selection correlations. Mann-Whitney U tests compared experimental values against null distributions. Chi-square tests for trend assessed outcome fraction shifts across nutrient conditions. Fisher’s exact test compared categorical outcomes between conditions. Linear regression quantified predictability of coalescence outcomes from dominant-species competition (R^2^ reported). For figures requiring error bars, the mean and s.e.m. are presented, with specific test details provided in each legend. Full statistical methods are provided in Supplementary Methods.

### Data Availability

Isolates and communities are available upon request. All data are available in the Supplementary Information and via Dryad at http://datadryad.org/share/LINK_NOT_FOR_PUBLICATION/kQACU7LCmQclVZfGZk0bS5ZPUVL_grhwah2zvFY4m9s.

### Code Availability

All codes used for simulation and analysis are available via GitHub at https://github.com/Jinyeop3110/interspecies-interaction-derive-Community-Level-Selection.

## Acknowledgements

J.G. acknowledges funding support from the Schmidt Polymath Award and the Sloan Foundation.

## Author Contributions

J.Y.S., J.H. and J.G. conceived the study. J.Y.S. and J.H. performed the experiments and the theoretical modeling. J.Y.S. analyzed the data. All authors wrote and edited the manuscript.

## Additional Information

Supplementary Information is available for this paper.

## References

[1] Clements, F. E. Plant succession: An analysis of the development of vegetation. Carnegie Institution of Washington Publication 242, 1–512 (1916).

[2] Clements, F. E. Nature and structure of the climax. Journal of Ecology 24, 252–284 (1936).

[3] Odum, E. P. The strategy of ecosystem development. Science 164, 262–270 (1969).

[4] Lovelock, J. E. Gaia: A New Look at Life on Earth (Oxford University Press, Oxford, 1979).

[5] Wilson, D. S. & Sober, E. Reviving the superorganism. Journal of Theoretical Biology 136, 337–356 (1989).

[6] Gleason, H. A. The individualistic concept of the plant association. The American Midland Naturalist 21, 92–110 (1939).

[7] Cain, S. A. Characteristics of natural areas and factors in their development. Ecological Monographs 17, 185–200 (1947).

[8] Mason, H. L. Evolution of certain floristic associations in western north america. Ecological Monographs 17, 201–210 (1947).

[9] Whittaker, R. H. Gradient analysis of vegetation. Biological Reviews 42, 207–264 (1967).

[10] Rillig, M. C. et al. Interchange of entire communities: Microbial community coalescence. Trends in Ecology & Evolution 30, 470–476 (2015).

[11] Lechón-Alonso, P., Clegg, T., Cook, J., Smith, T. P. & Pawar, S. The role of competition versus cooperation in microbial community coalescence. PLoS Computational Biology 17, e1009584 (2021).

[12] Huet, S. et al. Experimental community coalescence sheds light on microbial interactions in soil and restores impaired functions. Microbiome 11, 42 (2023).

[13] Bresciani, L., Custer, G. F., Koslicki, D. & Dini-Andreote, F. Interplay of ecological processes modulates microbial community reassembly following coalescence. The ISME Journal 19, wraf041 (2025).

[14] Liu, X. & Salles, J. F. Drivers and consequences of microbial community coalescence. The ISME Journal 18, wrae179 (2024).

[15] Sarkar, A. et al. Microbial transmission in the social microbiome and host health and disease. Cell 187, 17–43 (2024).

[16] Xiao, Y., Angulo, M. T., Lao, S., Weiss, S. T. & Liu, Y.-Y. An ecological framework to understand the efficacy of fecal microbiota transplantation. Nature Communications 11, 3329 (2020).

[17] Gupta, S., Allen-Vercoe, E., Bhattacharya, A. & Petrof, E. O. Fecal microbiota transplantation: in perspective. Therapeutic Advances in Gastroenterology 9, 229–239 (2016).

[18] Gilpin, M. Community-level competition: Asymmetrical dominance. Proceedings of the National Academy of Sciences 91, 3252–3254 (1994).

[19] Debray, R. et al. Priority effects in microbiome assembly. Nature Reviews Microbiology 20, 109–121 (2022).

[20] Venturelli, O. S. et al. Deciphering microbial interactions in synthetic human gut microbiome communities. Molecular Systems Biology 14, e8157 (2018).

[21] Maynard, D. S., Miller, Z. R. & Allesina, S. Predicting coexistence in experimental ecological communities. Nature Ecology & Evolution 4, 91–100 (2020).

[22] Tikhonov, M. Community-level cohesion without cooperation. eLife 5, e15747 (2016).

[23] Xie, L., Yuan, A. E. & Shou, W. Simulations reveal challenges to artificial community selection and possible strategies for success. PLOS Biology 17, e3000295 (2019).

[24] Xie, L. & Shou, W. Steering ecological-evolutionary dynamics to improve artificial selection of microbial communities. Nature Communications 12, 6799 (2021).

[25] Diaz-Colunga, J. et al. Top-down and bottom-up cohesiveness in microbial community coalescence. Proceedings of the National Academy of Sciences 119, e2111261119 (2022).

[26] Vermeij, G. J. When biotas meet: Understanding biotic interchange. Science 253, 1099–1104 (1991).

[27] Brochet, S. et al. Niche partitioning facilitates coexistence of closely related honey bee gut bacteria. eLife 10, e68583 (2021).

[28] Benton, M. J. & Emerson, B. C. How did life become so diverse? the dynamics of diversification according to the fossil record and molecular phylogenetics. Palaeontology 50, 23–40 (2007).

[29] Van der Gucht, K. et al. The power of species sorting: Local factors drive bacterial community composition over a wide range of spatial scales. Proceedings of the National Academy of Sciences 104, 20404–20409 (2007).

[30] Goldman, D. A. et al. Competition for shared resources increases dependence on initial population size during coalescence of gut microbial communities. Proceedings of the National Academy of Sciences 122, e2322440122 (2025).

[31] Walton, S. J. et al. Community coalescence reveals strong selection and coexistence within species in complex microbial communities. bioRxiv (2025).

[32] May, R. M. Will a large complex system be stable? Nature 238, 413–414 (1972).

[33] Grilli, J. et al. Feasibility and coexistence of large ecological communities. Nature Communications 8, 14389 (2017).

[34] Hu, J., Amor, D. R., Barbier, M., Bunin, G. & Gore, J. Emergent phases of ecological diversity and dynamics mapped in microcosms. Science 378, 85–89 (2022).

[35] Hu, J., Barbier, M., Bunin, G. & Gore, J. Collective dynamical regimes predict invasion success and impacts in microbial communities. Nature Ecology & Evolution 9, 406–416 (2025).

[36] Ratzke, C., Barrere, J. & Gore, J. Strength of species interactions determines biodiversity and stability in microbial communities. Nature Ecology & Evolution 4, 376–383 (2020).

[37] Castledine, M., Sierocinski, P., Padfield, D. & Buckling, A. Community coalescence: An eco-evolutionary perspective. Philosophical Transactions of the Royal Society B: Biological Sciences 375, 20190252 (2020).

[38] Ratzke, C. & Gore, J. Modifying and reacting to the environmental ph can drive bacterial interactions. PLOS Biology 16, e2004248 (2018).

[39] Nadell, C. D., Drescher, K. & Foster, K. R. Spatial structure, cooperation and competition in biofilms. Nature Reviews Microbiology 14, 589–600 (2016).

[40] Marsland III, R. et al. Available energy fluxes drive a transition in the diversity, stability, and functional structure of microbial communities. PLoS Computational Biology 15, e1006793 (2019).

[41] Allesina, S. & Tang, S. Stability criteria for complex ecosystems. Nature 483, 205–208 (2012).

[42] Kurkjian, H. M., Akbari, M. J. & Momeni, B. The impact of interactions on invasion and colonization resistance in microbial communities. PLOS Computational Biology 17, e1008643 (2021).

[43] Fukami, T. Historical contingency in community assembly: Integrating niches, species pools, and priority effects. Annual Review of Ecology, Evolution, and Systematics 46, 1–23 (2015).

[44] Rillig, M. C. & Mansour, I. Microbial ecology: Community coalescence stirs things up. Current Biology 27, R1280–R1282 (2017).

[45] Tropini, C., Earle, K. A., Huang, K. C. & Sonnenburg, J. L. The gut microbiome: Connecting spatial organization to function. Cell Host & Microbe 21, 433–442 (2017).

[46] Smillie, C. S. et al. Strain tracking reveals the determinants of bacterial engraftment in the human gut following fecal microbiota transplantation. Cell Host & Microbe 23, 229–240.e5 (2018).

[47] Rocca, J. D., Muscarella, M. E., Peralta, A. L., Izabel-Shen, D. & Simonin, M. Guided by microbes: Applying community coalescence principles for predictive microbiome engineering. mSystems 6, e00538–21 (2021).

[48] Callahan, B. J. et al. Dada2: High-resolution sample inference from illumina amplicon data. Nature Methods 13, 581–583 (2016).

[49] Quast, C. et al. The silva ribosomal rna gene database project: improved data processing and web-based tools. Nucleic Acids Research 41, D590–D596 (2013).

