## Supplementary Materials for "Interspecies Interactions Drive Community-Level Selection in Microbial Coalescence"

#### Supplementary Methods

##### Microbial Strain Library, Media, and Culture

We obtained a library of 54 bacterial isolates derived from soil, tree surface, and flower stamen environments collected in Cambridge, MA, USA. Each isolate was purified by serial streaking and stored as glycerol stocks (25% glycerol, −80°C). Phylogenetic identities were determined by full-length 16S rRNA gene sequencing and V4 region amplicon sequencing with BLAST alignment to the SILVA 138 database; the 54 isolates corresponded to 50 unique amplicon sequence variants (ASVs) spanning 29 families across three phyla (Proteobacteria, Firmicutes, and Bacteroidota) (Extended Data Fig. 1; Supplementary Fig. 1). Monoculture growth characteristics were measured for all 54 isolates prior to community assembly experiments. Growth yield ranged from $OD_{600} = 0.1$ to 1.2 after 24 h, and final pH in unbuffered medium ranged from 4.5 to 8.5, indicating substantial phenotypic diversity in growth rate and environmental modification potential (Supplementary Figs. 8, 20).

The culture medium (CM) for isolates was composed of 1 g $L^{-1}$ yeast extract and 1 g $L^{-1}$ soytone (both from Becton Dickinson), 10 mM sodium phosphate buffer adjusted to pH 6.5, 0.1 mM $CaCl_2$, 2 mM $MgCl_2$, 4 mg $L^{-1}$ $NiSO_4$, and 50 mg $L^{-1}$ $MnCl_2$. All media were filter sterilized using bottle-top filtration units (VWR), and all chemicals were purchased from Sigma–Aldrich unless otherwise noted.

Our coalescence experiments were conducted in three media conditions with varying nutrient supplementation, which produce different strengths of pairwise interactions: (1) Nutr−, the culture medium without additional carbon or nitrogen supplementation; (2) Base, CM supplemented with 5 g $L^{-1}$ glucose and 4 g $L^{-1}$ urea; and (3) Nutr+, CM supplemented with 20 g $L^{-1}$ glucose and 16 g $L^{-1}$ urea. Both parental community assembly and coalescence experiments were performed in the same media condition throughout each experimental series.

Both monocultures and communities were grown in 300 $\mu$L volumes in 96-well deep-well plates (Deepwell plate 96/1000 $\mu$L; Eppendorf) covered with AeraSeal adhesive sealing films (Excel Scientific). Plates were incubated aerobically at 25°C and shaken at 800 r.p.m. on Titramax shakers (Heidolph). To minimize pre-adaptation, all isolates from frozen stocks were preconditioned for two serial growth–dilution cycles prior to assembling parental communities.

##### Construction of Parental Communities and Community Coalescence

Parental communities were constructed at three richness levels: 6, 12, or 24 members (P6, P12, P24), with 30 total communities (9 × P6, 9 × P12, 12 × P24). For P24 communities, 12 partially overlapping communities were assembled because only two non-overlapping sets of 24 species can be drawn from 54 isolates. Coalescence pairs were composed systematically: P6 included 14 pairs, P12 included 27 pairs, and P24 included 6 non-overlapping pairs, resulting in 47 total coalescence pairs for synthetic communities. Some coalescence events were excluded due to sequencing failures. After seven daily growth–dilution cycles, parental communities in Base medium retained a mean species survival ratio of $74 \pm 2\%$ (mean ± s.e.m.) of their initial species pool, indicating that Base medium supports high species coexistence under moderate interaction strength.

Coalescence experiments were conducted by mixing two pre-stabilized parental communities (A and B) at equal volume ratio (1:1) after seven daily dilution cycles. The resulting mixtures were cultured for another seven serial transfers (×30 dilutions every 24 h) under identical media conditions to reach new steady states (Fig. 1c). Each of the 47 coalescence pairs was performed with two biological replicates, yielding 94 total coalescence events; 11 events were excluded due to sequencing failures or contamination, resulting in 83 coalescence events for synthetic community analyses.

##### 16S rRNA Sequencing and Data Processing

Genomic DNA was extracted using QIAGEN DNeasy PowerSoil HTP 96 Kit following the manufacturer's protocol, with bead-beating for cell lysis. The V4 region of the 16S rRNA gene was amplified using 515F/806R primers containing Illumina overhang adapters and sequenced on an Illumina MiSeq platform (2×250 bp). Raw reads were demultiplexed and denoised using QIIME2 and DADA2, generating amplicon sequence variants (ASVs). ASVs were taxonomically classified against the SILVA v138 database.

Relative abundances were normalized by total read depth. ASV tables were filtered to retain only taxa representing ≥0.1% relative abundance in any sample, which corresponds to the 0.1% extinction threshold used in simulations. The phylogenetic tree (Supplementary Fig. 1) was constructed using Simple Phylogeny by EMBL's European Bioinformatics Institute[1].

**Similarity-Based Classification of Coalescence Outcomes**

To characterize coalescence outcomes, we quantified the compositional similarity between the post-coalescence community and each parental community. Each community's species abundance profile was normalized to unit length using $L_2$ normalization (i.e., dividing by the square root of the sum of squared abundances, so that the resulting vector has unit Euclidean length). We then computed the cosine similarity (dot product of normalized vectors) between the coalesced community C and each parent A and B. These two similarity scores place each coalescence outcome in a two-dimensional similarity space, where the position reflects how strongly the coalesced community resembles each parental community.

Because parental communities may share species (overlapping ASVs), we used a linear decomposition approach to determine how much of the coalesced community's composition can be attributed to each parental community. We computed contribution coefficients $(u, v)$ representing the weighted combination of parental compositions that best reconstructs the coalesced community. When parental communities share no species, these coefficients equal the cosine similarities. Negative coefficients were set to zero since negative contributions are biologically meaningless. The residual (the portion of composition not explained by either parental community) quantifies novel restructuring.

From these coefficients, we derived two classification metrics:

Retention magnitude ($x$): The overall contribution from both parental communities, calculated as $x = \sqrt{u^2 + v^2}$. Values close to 1 indicate high retention of parental composition; values close to 0 indicate extensive restructuring.

Parental Dominance Index (PDI): A measure of selection preference toward one parent over the other, ranging from 0 (complete dominance by parent B) through 0.5 (equal contributions) to 1 (complete dominance by parent A). PDI is computed from the angular position in similarity space relative to the diagonal where both parental communities contribute equally:

$$\text{PDI} = \frac{2}{\pi} \arctan\left(\frac{u}{v}\right) \tag{1}$$

where $u$ and $v$ are the contribution coefficients for parents A and B, respectively. When $u \gg v$, PDI approaches 1 (parent A dominance); when $v \gg u$, PDI approaches 0 (parent B dominance); when $u = v$, PDI equals 0.5 (equal contributions).

Based on these metrics, each coalescence event was classified into one of three outcome categories. Restructuring occurs when $x^2 \leq 0.5$, meaning less than half of the coalesced community composition is explained by the parental communities, indicating substantial ecological reorganization. Mixture occurs when $x^2 > 0.5$ and PDI is near 0.5 (i.e., $0.25 \leq \text{PDI} \leq 0.75$), indicating high retention with balanced contributions from both parental communities and coexistence of parental lineages. Dominance occurs when $x^2 > 0.5$ and PDI is near 0 or 1 (i.e., PDI $< 0.25$ or PDI $> 0.75$), indicating high retention but with one parental community dominating, reflecting selection for one parental community over the other.

We chose the classification thresholds based on two considerations. First, the retention threshold $x^2 = 0.5$ represents the midpoint where parental communities explain exactly half of the offspring composition; values below this indicate that novel species combinations dominate over parental contributions. Second, the PDI thresholds of 0.25 and 0.75 correspond to contribution ratios of approximately 3:1 between parental communities, a level of asymmetry that represents clear dominance of one parent over the other while allowing for minor contributions from the subordinate parent.

##### Lotka–Volterra Simulations

We modeled species-abundance dynamics with the generalized Lotka–Volterra (gLV) system:

$$\frac{dn_i}{dt} = n_i \left( r_i - \sum_{j=1}^{S} \alpha_{ij} n_j \right) \tag{2}$$

where $n_i$ is the abundance of species $i$, $r_i$ is its intrinsic growth rate, $S$ is the total number of species, and $\alpha_{ij}$ is the interaction coefficient describing how strongly species $j$ inhibits the growth of species $i$ (with self-regulation $\alpha_{ii} = 1$). To isolate the role of interaction strength, we set all growth rates $r_i = 1$. Off-diagonal interaction coefficients ($i \neq j$) were drawn randomly from a uniform distribution $U[0, 2\mu]$ with mean $\mu$, which controls the average strength of interspecific competition.

Each simulation used a pool of $N = 54$ species partitioned into four non-overlapping parental communities of 12 species. For each replicate, we randomly permuted the 54 species and assigned them sequentially (1–12, 13–24, 25–36, 37–48) to form the four communities, ensuring no species were shared across parental communities, analogous to the experimental design. A fresh interaction matrix was independently sampled for every replicate so that coalescence dynamics were explored across diverse ecological contexts rather than a single fixed species pool.

For single-community assembly, we numerically integrated the gLV equations from random initial conditions $N_i(0) \sim \text{Unif}(0, 0.1)$ to ecological steady state. We solved the ODEs with the adaptive Runge–Kutta method RK23 implemented in `scipy.integrate.solve_ivp`, integrating from $t = 0$ to $t = 5000$, which was sufficient for all communities to equilibrate. Following integration, species with relative abundances below an extinction threshold of 0.1% were set to zero, yielding four parental-community steady states per replicate.

Community coalescence was simulated by mixing pre-equilibrated parental communities at equal proportions, mirroring the experimental 1:1 volume protocol. For each of the $\binom{4}{2} = 6$ parental community pairs, we retrieved the steady-state abundance vectors $\mathbf{y}^{(1)}$ and $\mathbf{y}^{(2)}$, formed the equal mixture $\mathbf{y}_{\text{mix}} = (\mathbf{y}^{(1)} + \mathbf{y}^{(2)})/2$, and zeroed species falling below the extinction threshold in this initial mixture. We then re-integrated the gLV system from $\mathbf{y}_{\text{mix}}$ using the same interaction matrix over $t \in [0, 5000]$ until a new steady state was reached, and reapplied the 0.1% threshold to the final abundances. This procedure captures the ecological relaxation after mixing and allows competitive exclusion, coexistence, or the emergence of previously rare species to arise from the underlying interaction structure.

##### Pairwise Invasion Assays

To empirically estimate interaction strength, we performed pairwise invasion experiments among the 12 most abundant isolates. Isolate pairs were mixed at a 95:5 ratio (resident:invader) by colony count and propagated in Nutr−, Base, and Nutr+ media under daily dilution (×30) for 7 cycles. Final compositions were determined by colony counting on agar plates. Competitive outcomes were scored as coexistence, exclusion, or bistability based on final relative abundances (coexistence if both >10%; exclusion if one <1%). The fraction of failed invasions was used as a proxy for mean interaction strength under each nutrient condition, as higher failed invasion frequency indicates a greater proportion of pairwise interaction coefficients exceeding 1 in the gLV framework.

##### Natural Sample-Derived Communities

Six environmental samples were collected from diverse microhabitats in Cambridge, MA, USA (soil, compost, decomposing organic matter). Each sample was homogenized and inoculated into culture medium, then subjected to seven serial growth-dilution cycles (×30 dilution every 24 h) to establish stable communities. After stabilization, 15 pairwise coalescence events (each with two biological replicates, $n = 30$ per condition) were performed across all three media conditions following the same protocol as synthetic communities. Community composition was characterized via 16S rRNA amplicon sequencing.

##### Statistical Analyses

All analyses were performed in Python 3.11 using NumPy, pandas, and scikit-learn, with statistical significance defined at $p < 0.05$ unless otherwise noted.

- Paired $t$-tests were used to compare mean interaction strengths before and after community assembly.
- Permutation tests (1,000 permutations) were used to assess whether pairwise selection correlations differed between same-parent and cross-parent species pairs. The null distribution was generated by shuffling species origin labels within each coalescence event.
- Mann-Whitney U tests were used to compare experimental Dominance values against null model distributions.
- Linear regression was used to quantify the predictability of coalescence outcomes from dominant-species pairwise competition, with $R^2$ reported as the coefficient of determination.
- $\chi^2$ tests were used to compare observed vs. expected counts of Dominance, Mixture, and Restructuring events.
- Mixed-effects models (lme4 package in R) were used to control for parental community richness and community identity effects when quantifying nutrient dependence.

##### Optical Density Measurements and Monoculture Characterization

Optical density (OD) was measured at 600 nm using a plate reader (BioTek Synergy H1) to monitor community growth and standardize inoculation densities. For community assembly, parental communities were normalized to equivalent $OD_{600}$ prior to coalescence mixing to ensure equal biomass contributions from each parent. Post-coalescence samples were measured at each serial dilution cycle to track growth dynamics. All OD measurements were blanked against sterile media controls.

To characterize the phenotypic diversity of isolates prior to community assembly, we measured growth rate and pH modification potential for each of the 54 bacterial isolates in monoculture. $OD_{600}$ was recorded at 24-hour intervals during a 7-day serial dilution regime ($\times 30$ dilution every 24 h) in the culture medium (CM), and growth rate was estimated from the exponential phase. Final pH was measured using a microplate pH meter after 24 h of growth in unbuffered medium to assess each isolate's capacity to acidify or alkalinize the environment. The 54 isolates exhibited broad variation in both growth yield ($OD_{600}$ range: 0.1–1.2) and pH modification potential (final pH range: 4.5–8.5), providing the phenotypic diversity necessary for diverse coalescence outcomes (Supplementary Figs. 8, 20).

##### Sensitivity Analysis

To assess the robustness of our classification scheme, we performed sensitivity analyses on similarity metrics and simulation parameters.

Similarity Metric Robustness: We compared coalescence outcome classifications using five different similarity metrics: vector decomposition (primary method), Euclidean distance, Bray-Curtis dissimilarity, Jensen-Shannon divergence, and Jaccard index. Despite different mathematical formulations, all metrics yielded qualitatively consistent outcome distributions, with Dominance, Mixture, and Restructuring fractions varying by <15% across metrics (Extended Data Fig. 2).

Simulation Robustness: For Lotka–Volterra simulations, we tested alternative interaction coefficient distributions, including Gaussian (mean $\mu$, standard deviation $\mu/\sqrt{3}$, truncated at 0) and Gamma (mean $\mu$, shape parameter 3) distributions with matched variance $\mu^2/3$. The qualitative transition from Mixture-dominated to Dominance/Restructuring outcomes with increasing interaction strength was robust to distributional choice (Supplementary Fig. 4).

#### Supplementary Note 1: Null Models and Statistical Controls

A potential concern is that Dominance could arise from skewed abundance distributions rather than ecological interactions. To address this, we compared experimental Dominance values against two null models.

In Null Model 1 (abundance-weighted random selection), species survival probability is proportional to abundance in the combined parental pool, testing whether high-abundance species simply survive regardless of parental origin. In Null Model 2 (shuffled abundance), abundances are randomly permuted among species within each parent before neutral mixing, testing whether the overall skewness structure alone drives Dominance.

We generated 500 null offspring communities per model and calculated the Dominance metric. Experimental data showed mean Dominance of 0.698 ($n = 83$). Both null models produced significantly lower values: abundance-weighted null mean = 0.476 ($p < 10^{-18}$), shuffled null mean = 0.557 ($p < 10^{-11}$; Mann-Whitney U tests). These results indicate that abundance skewness alone cannot explain the observed patterns; ecological interactions beyond simple abundance-weighted survival determine which parental community dominates (Extended Data Fig. 3).

#### Supplementary Note 2: Assembly Effect Analysis

To understand the mechanistic basis of community-level selection, we compared coalescence outcomes to a "direct assembly" control. In coalescence, two parental communities are first assembled separately from the species pool, then mixed; in direct assembly, all species from both pools are mixed simultaneously without prior assembly history.

During assembly, species compete and some go extinct. Those that survive to coexistence tend to have weaker mutual interactions on average. We quantified this assembly effect by measuring the mean pairwise interaction strength $\langle \alpha_{ij} \rangle$ among species at different stages (Supplementary Fig. 18). Across all tested interaction strengths ($\mu = 0$ to 1.2), post-assembly communities exhibited substantially reduced mean interaction strength compared to the pre-assembly pool. This reduction becomes more pronounced at higher $\mu$, where stronger competition drives more species to extinction, leaving only those with weaker mutual interactions. The assembly process thus creates internal coherence within each parental community: species that survived together share compatible (less competitive) interactions.

We hypothesized that this shared history would increase Dominance probability compared to direct assembly. Simulations supported this prediction (Extended Data Fig. 7): across interaction strengths $\mu = 0.2$ to 1.0, coalescence consistently produced higher Dominance fractions than direct assembly. At $\mu = 0.2$, coalescence yielded 37% Dominance versus 12% for direct assembly; at $\mu = 0.6$, 59% versus 20%; at $\mu = 1.0$, 65% versus 22%. Correspondingly, direct assembly showed higher Restructuring rates, as expected when species from different pools have not been pre-filtered for compatibility.

Experimental results corroborate these findings (Supplementary Fig. 19). Under Base and Nutr+ conditions, coalescence showed elevated Dominance rates compared to direct assembly, consistent with simulation predictions. The Nutr− condition showed more variable results, likely due to weaker competitive interactions in nutrient-limited environments.

#### Supplementary Note 3: Simulation Robustness

To test the robustness of our simulation results to the choice of interaction coefficient distribution, we compared three distributions with matched mean interaction strength $\mu$: uniform ranging from 0 to $2\mu$ (primary analysis), Gaussian with mean $\mu$ and standard deviation $\mu/\sqrt{3}$ (truncated at 0), and Gamma with mean $\mu$ and shape parameter 3. All three distributions were parameterized to have the same mean $\mu$ and matched variance $\mu^2/3$ (for the uniform distribution $U[0, 2\mu]$, var $= (2\mu)^2/12 = \mu^2/3$). The qualitative transition from Mixture-dominated to Dominance/Restructuring outcomes with increasing interaction strength was robust across all tested distributions (Supplementary Fig. 4).

We also tested whether community size (number of species per parental community) affects the frequency of Dominance outcomes. Simulations were performed with parental community sizes ranging from 4 to 48 species per community (Extended Data Fig. 5). The qualitative patterns were consistent across community sizes: Dominance frequency remained relatively stable ($\sim$14–26% at $\mu = 0.3$, $\sim$56–67% at $\mu = 0.6$, $\sim$69–78% at $\mu = 0.8$), while Mixture decreased and Restructuring increased with larger species pools. This pattern reflects increased opportunity for competitive exclusion cascades in larger communities, but the qualitative prevalence of Dominance at moderate-to-high interaction strengths is robust to community size.

#### Supplementary Note 4: Pairwise Selection Correlation

To quantify whether species from the same parental community exhibit correlated selection during coalescence, we developed a pairwise selection correlation metric. For each coalescence event, species were assigned to their parental community of origin (the parent with higher abundance if present in both). For each species pair, we evaluated whether their presence/absence patterns in the offspring were concordant (both present or both absent) or discordant (one survives, one extinct).

The concordance rate was converted to a correlation metric: $\rho = 2 \times \text{Concordance rate} - 1$, yielding values from $-1$ (all discordant) to $+1$ (all concordant). We computed $\rho_{\text{same}}$ for within-community pairs and $\rho_{\text{cross}}$ for cross-community pairs, then averaged across coalescence events. To test significance, we generated null distributions by shuffling species origin labels (1,000 permutations) and computed permutation $p$-values for $\Delta = \bar{\rho}_{\text{same}} - \bar{\rho}_{\text{cross}}$.

Positive $\bar{\rho}_{\text{same}}$ indicates species from the same parental community share fates (survive or go extinct together); negative $\bar{\rho}_{\text{cross}}$ indicates cross-community species have opposite fates. In both simulations (Extended Data Fig. 6a–d) and experiments (Extended Data Fig. 7), we observed $\bar{\rho}_{\text{same}} > 0$ and $\bar{\rho}_{\text{cross}} < 0$, with $\Delta$ increasing with interaction strength. At weak interactions ($\mu = 0.3$), both within-community and cross-community pairs show positive correlations with only a small difference ($\Delta = 0.05$; Extended Data Fig. 6a). As interaction strength increases, the difference becomes pronounced: at $\mu = 0.6$, within-community pairs show positive correlation while cross-community pairs become negative ($\Delta = 0.58$; Extended Data Fig. 6b); at $\mu = 0.8$, the separation is even larger ($\Delta = 0.94$; Extended Data Fig. 6c). This confirms that assembly history creates positive pairwise selection correlation among within-community species, providing the mechanistic basis for community-level selection.

#### Supplementary Note 5: Pairwise Invasion Experiments

To directly measure pairwise interactions between species, we performed reciprocal invasion assays using the 12 most abundant isolates (ranked by ASV counts across all communities). Each pair of isolates was tested in both directions: species A as resident with species B as invader, and vice versa. This reciprocal design allows us to assess competitive outcomes and detect potential bistability where the outcome depends on initial conditions.

Each invasion assay was initiated by mixing isolate pairs at a 95:5 ratio (resident:invader) by colony count from equal-biomass cultures. The co-cultures were then subjected to 7 cycles of daily dilution ($\times 30$ every 24 h) under the same growth conditions as community experiments (800 r.p.m. shaking at 25°C in 96-well deep-well plates). Final compositions were determined by colony counting on agar plates after the seventh dilution cycle. Invasion outcomes were classified as coexistence (both species maintained >10% relative abundance), exclusion (one species fell below 1% relative abundance), or bistability (outcome depended on which species started as resident).

Invasion assays were performed across all three nutrient conditions: Nutr−, Base, and Nutr+. The frequency of competitive exclusion increased with nutrient concentration, consistent with the coalescence outcome patterns observed in community experiments. Under Nutr−, more pairs achieved stable coexistence, while Nutr+ conditions led to more frequent exclusion of one species by the other.

#### Extended Data Figures

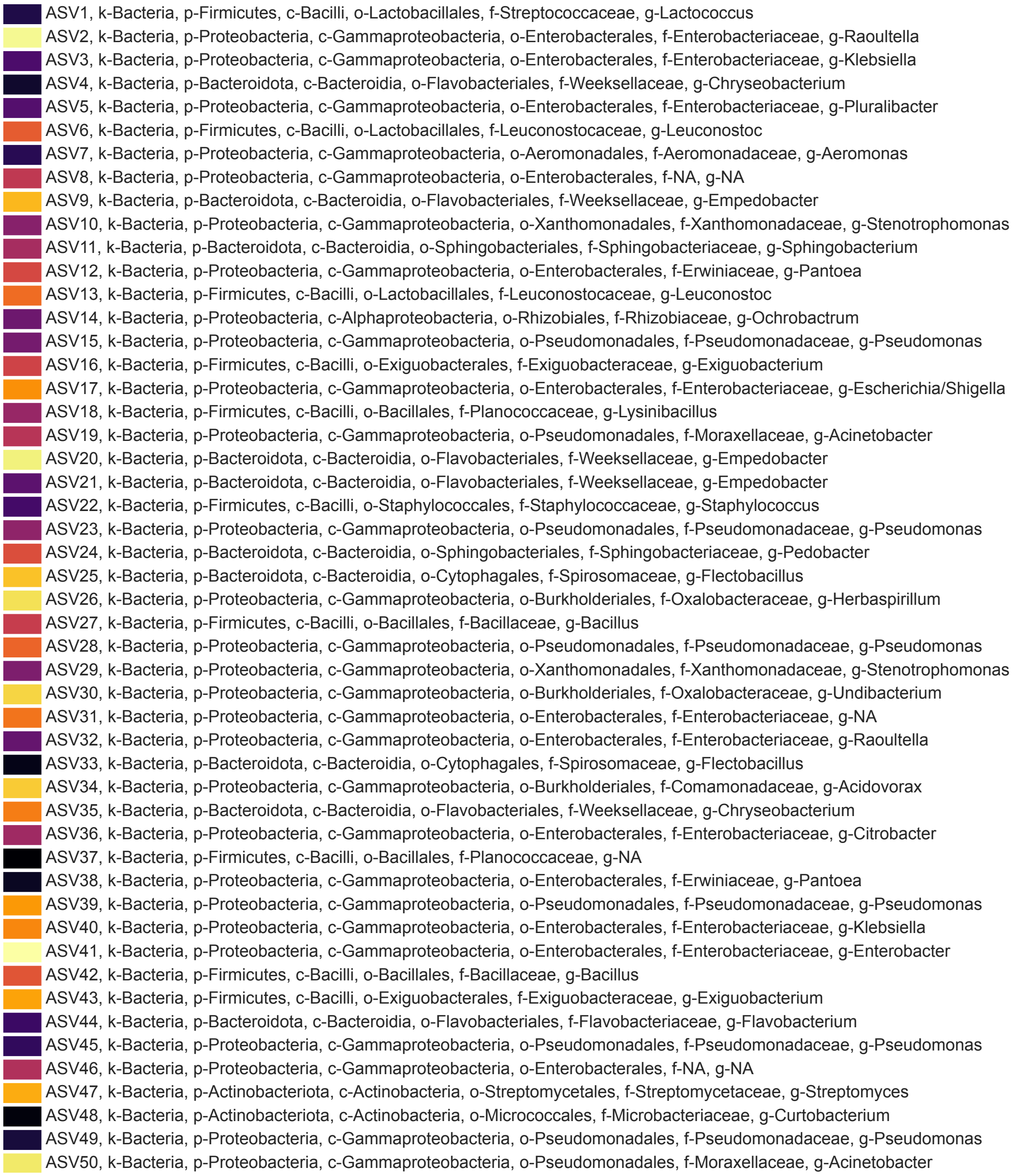


**Extended Data Fig. 1. Taxonomic classification of bacterial isolates.** Table showing the 54 bacterial isolates used in the experiment, of which 4 pairs share identical ASV sequences. Each row shows the ASV identifier and its full taxonomic classification (Kingdom, Phylum, Class, Order, Family, Genus). Colors match those used in pie charts and composition figures throughout the manuscript.

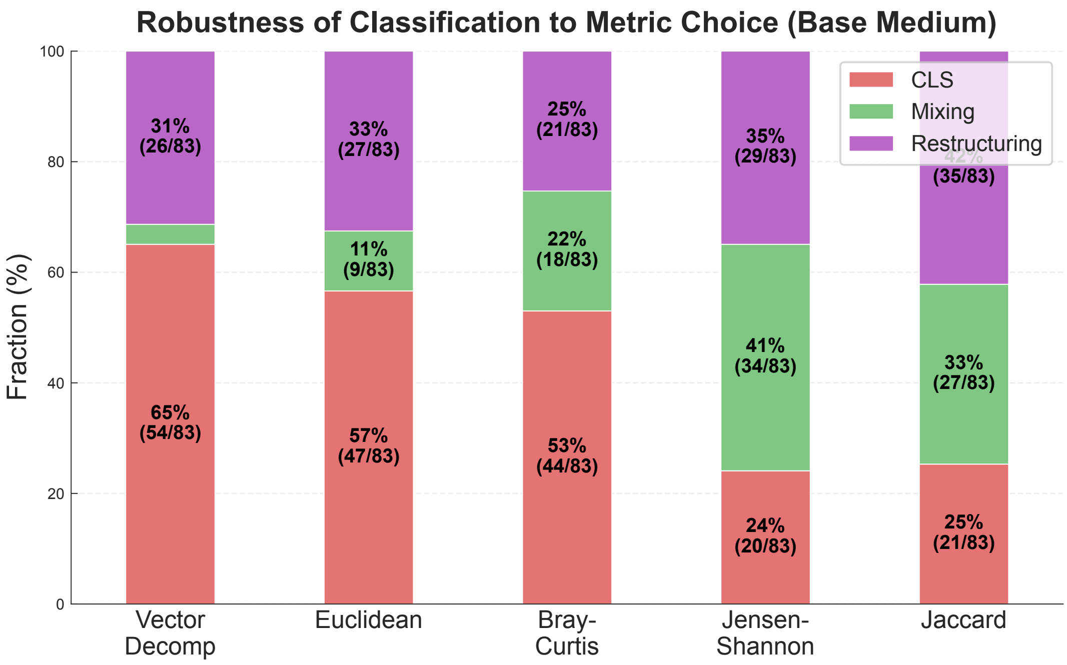


**Extended Data Fig. 2. Robustness of outcome classification to metric choice.** Stacked bar plot showing the distribution of coalescence outcomes (Dominance, Mixture, Restructuring) in Base medium using five different similarity metrics: Vector Decomposition, Euclidean distance, Bray-Curtis dissimilarity, Jensen-Shannon divergence, and Jaccard index. Percentages and counts (n/total) are shown for each category. All metrics produce qualitatively similar outcome distributions, demonstrating that the prevalence of Dominance is robust to the choice of similarity metric.

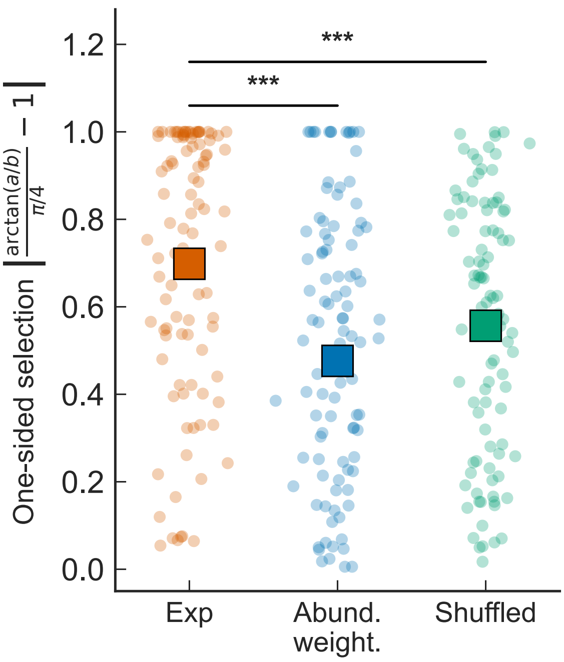


**Extended Data Fig. 3. Testing whether skewed parental abundance distributions explain Dominance.** Scatter plots show 100 randomly sampled data points with mean ± s.e.m. (squares with error bars) for experimental Dominance values compared against two null models: (1) abundance-weighted random selection, where species survival probability is proportional to their abundance in the combined parental pool, and (2) shuffled abundance, where abundances are randomly permuted among species within each parental community before neutral mixing. Both null models produce significantly lower Dominance than experimental data (Mann-Whitney U test, $p < 0.001$ for both comparisons), indicating that skewed abundance distributions alone do not fully account for the observed asymmetric outcomes.

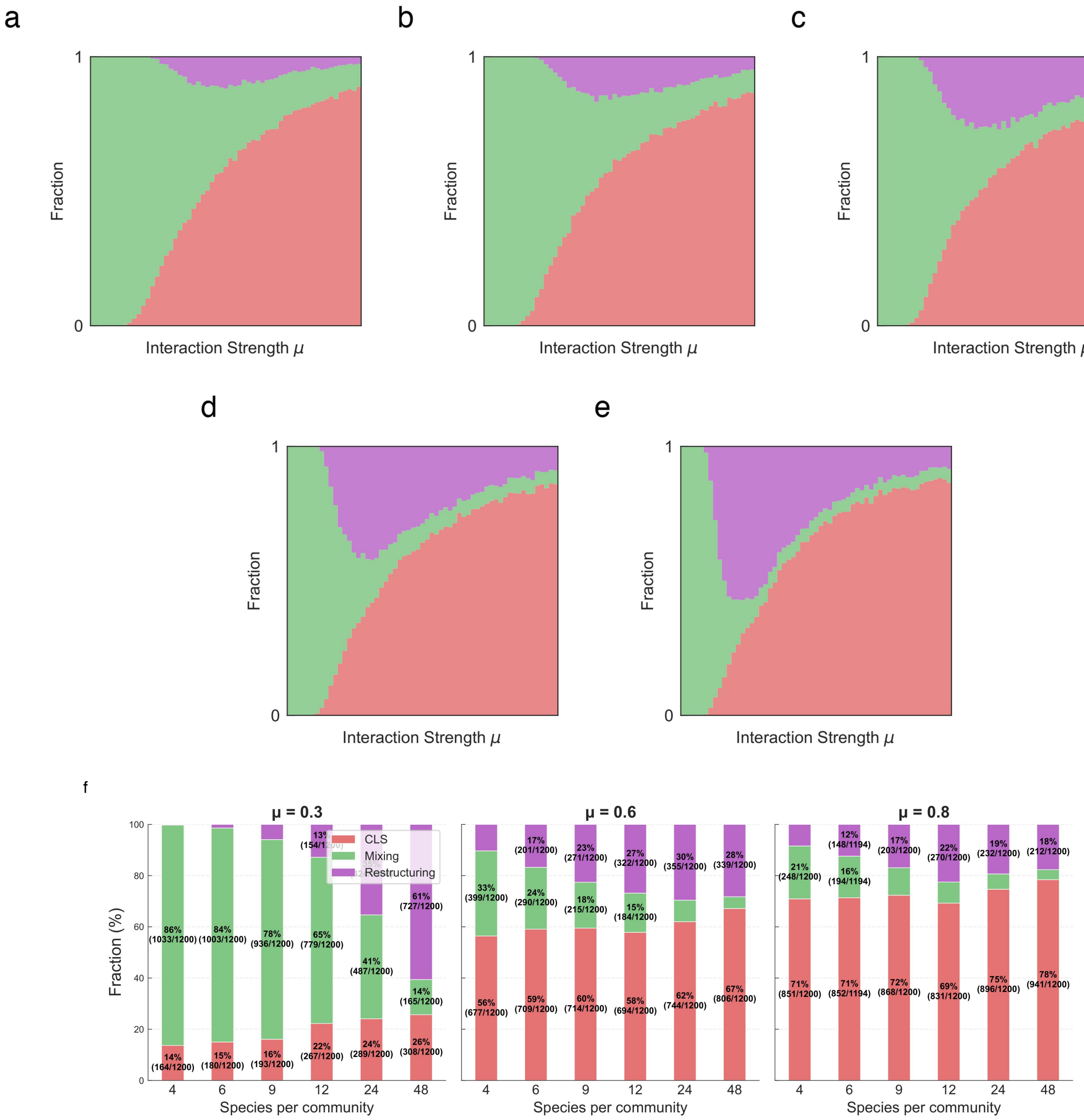


**Extended Data Fig. 4. Effect of community size on coalescence outcomes. (a–e)** Phase diagrams showing coalescence outcomes in similarity space for parental communities with 4, 6, 12, 24, and 48 species per community at $\mu = 0.6$. The qualitative pattern of Dominance-dominated outcomes is consistent across community sizes. **(f)** Stacked bar plots showing outcome fractions (Dominance, Mixture, Restructuring) across community sizes at three interaction strengths ($\mu = 0.3, 0.6, 0.8$). Dominance frequency remains relatively stable across community sizes, while Mixture decreases and Restructuring increases with larger communities. Simulations used 200 replicates per condition with uniform interaction coefficients $\alpha_{ij} \sim U(0, 2\mu)$.

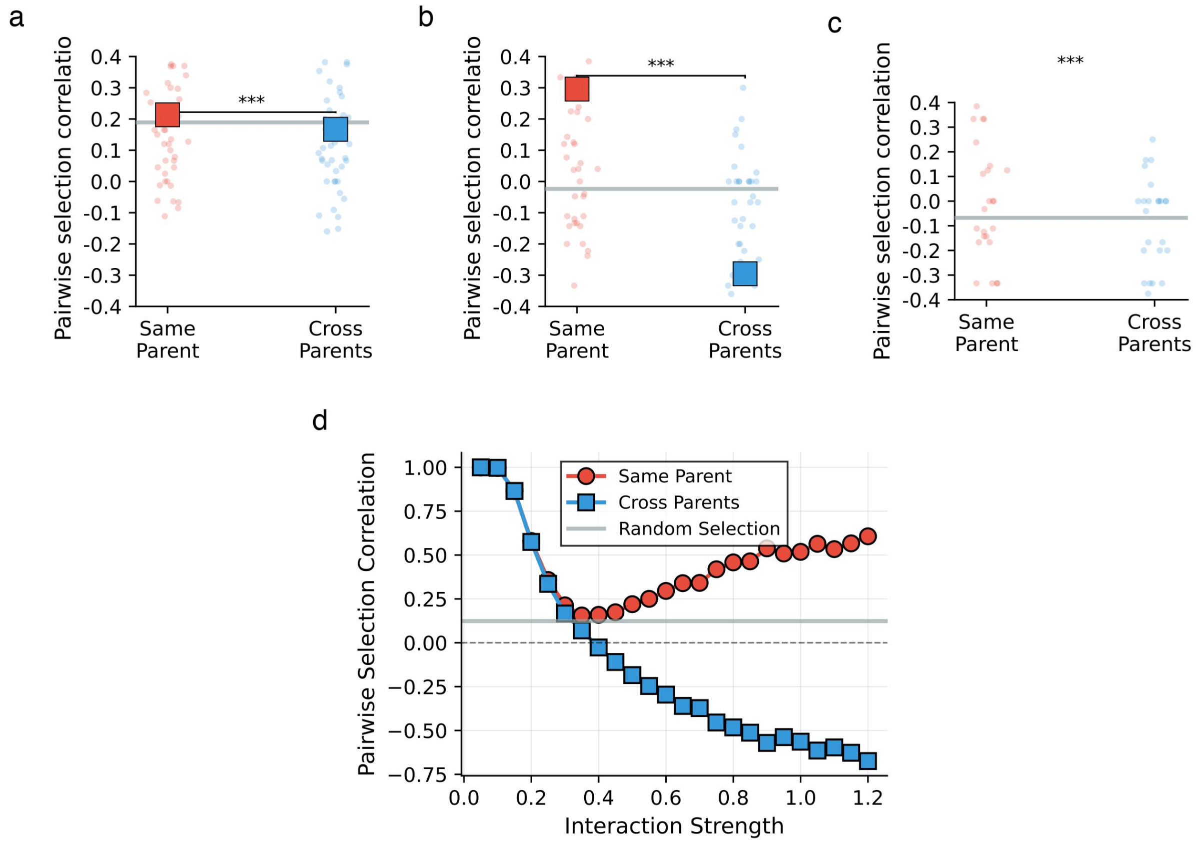


**Extended Data Fig. 5. Pairwise selection correlation increases with interaction strength in simulations. (a–c)** Pairwise selection correlation for species pairs from the same parental community (red, "Same parental community") versus cross-community pairs (blue, "Cross-community") at three representative interaction strengths. Gray horizontal lines indicate random selection baseline from null model. Individual dots show per-event correlations (50 stratified samples displayed); squares with error bars show mean ± s.e.m. At weak interactions ($\mu = 0.3$), both within-community and cross-community pairs show positive correlations with small difference ($\Delta = 0.05$). As interaction strength increases ($\mu = 0.6$, 0.8), within-community pairs maintain positive correlation while cross-community pairs become increasingly negative, indicating stronger community-level selection. **(d)** Mean pairwise selection correlation across all simulated coalescence events as a function of mean interaction strength $\mu$. Species pairs from the same parental community (red circles) show positive correlation that increases with $\mu$, while cross-community pairs (blue squares) show negative correlation that becomes more negative with $\mu$. Error bars represent standard error across $n = 1{,}200$ coalescence events per interaction strength (200 independent replicates × 6 pairwise coalescence events per replicate).

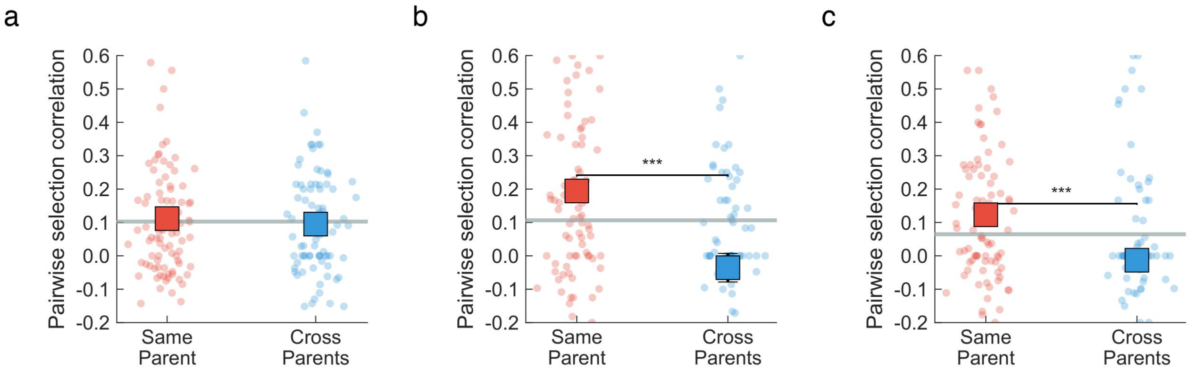


**Extended Data Fig. 6. Pairwise selection correlation in experimental coalescence across nutrient conditions.** Mean pairwise selection correlation for species pairs from the same parental community (red) versus cross-community pairs (blue). Gray horizontal line indicates random selection baseline. Individual dots show per-event correlations; squares with error bars show mean ± s.e.m. **(a)** In Nutr− medium, where interactions are weak, within-community and cross-community pairs show no significant difference in selection correlation, consistent with weak community-level selection. **(b)** In Base medium, intermediate differences emerge. **(c)** In Nutr+ medium, where interactions are strong, within-community species exhibit significantly higher selection correlation than cross-community pairs ($p < 0.001$), indicating positive pairwise selection correlation that underlies community-level selection.

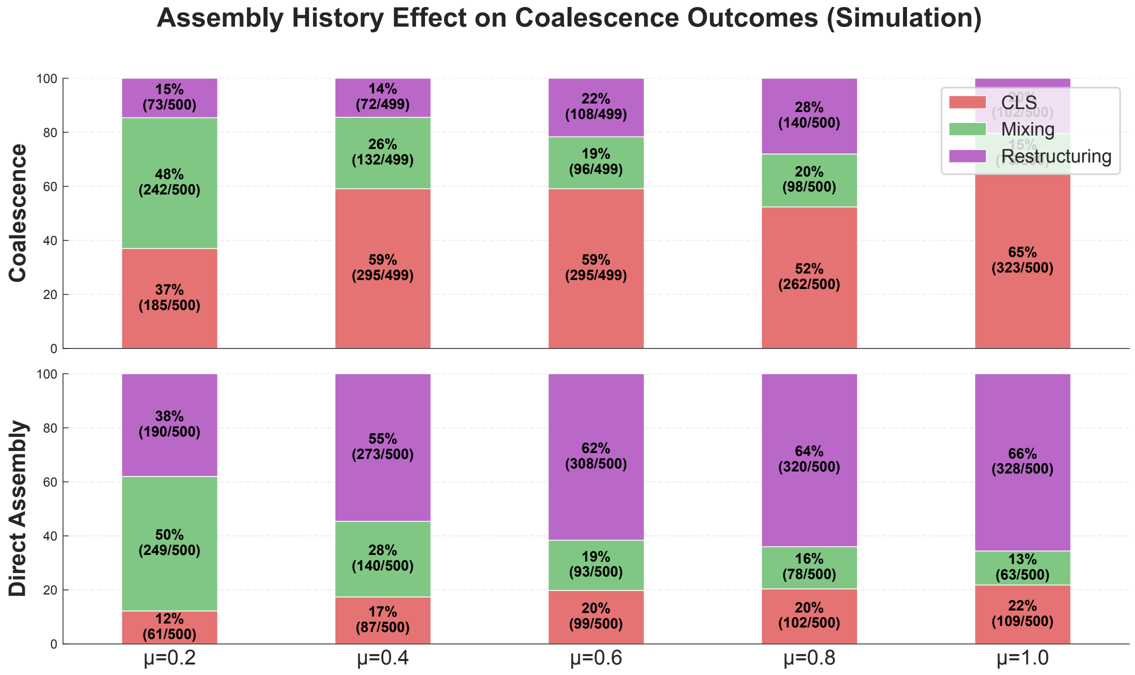


**Extended Data Fig. 7. Assembly history promotes Dominance in simulations.** Stacked bar plots comparing outcome distributions between coalescence (top row) and direct assembly (bottom row) across five interaction strengths ($\mu = 0.2, 0.4, 0.6, 0.8, 1.0$). Percentages and counts (n/total) are shown for each category. Coalescence consistently produces higher Dominance and lower Restructuring compared to direct assembly, demonstrating that assembly history creates correlated species retention.

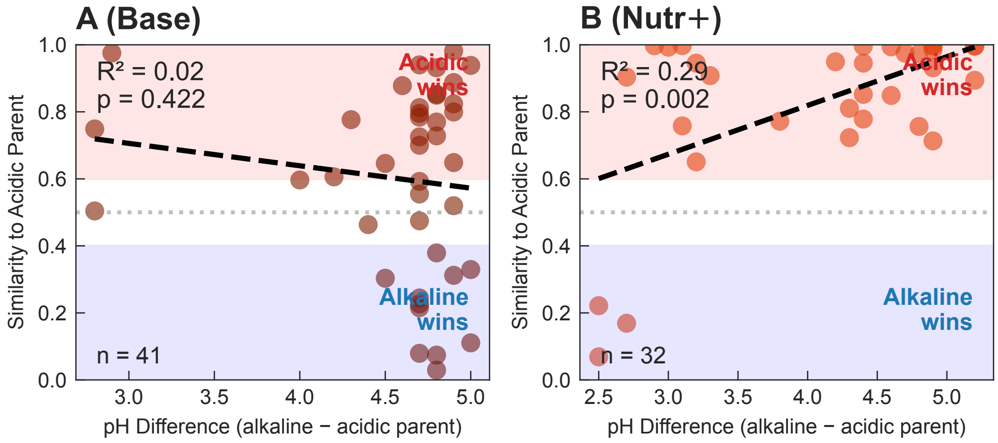


**Extended Data Fig. 8. pH difference between parental communities predicts coalescence outcome under strong interactions.** In coalescence events where one parental community is acidic (pH $< 6.5$) and the other alkaline (pH $> 7.5$), the pH difference predicts which community dominates under strong interactions. **(a)** Base medium ($n = 41$; $R^2 = 0.02$, $P = 0.42$, n.s.) and **(b)** Nutr+ medium ($n = 32$; $R^2 = 0.29$, $P = 0.002$). X-axis: pH difference (alkaline parental community pH $-$ acidic parental community pH). Y-axis: similarity of the coalesced community to the acidic parental community (PDI). Red shaded region indicates acidic community wins (PDI $> 0.6$); blue shaded region indicates alkaline community wins (PDI $<$ 0.4). The relationship is significant only in Nutr+, where amplified metabolic activity intensifies pH modification effects, providing mechanistic support for the top-down regime.

#### Supplementary Figures

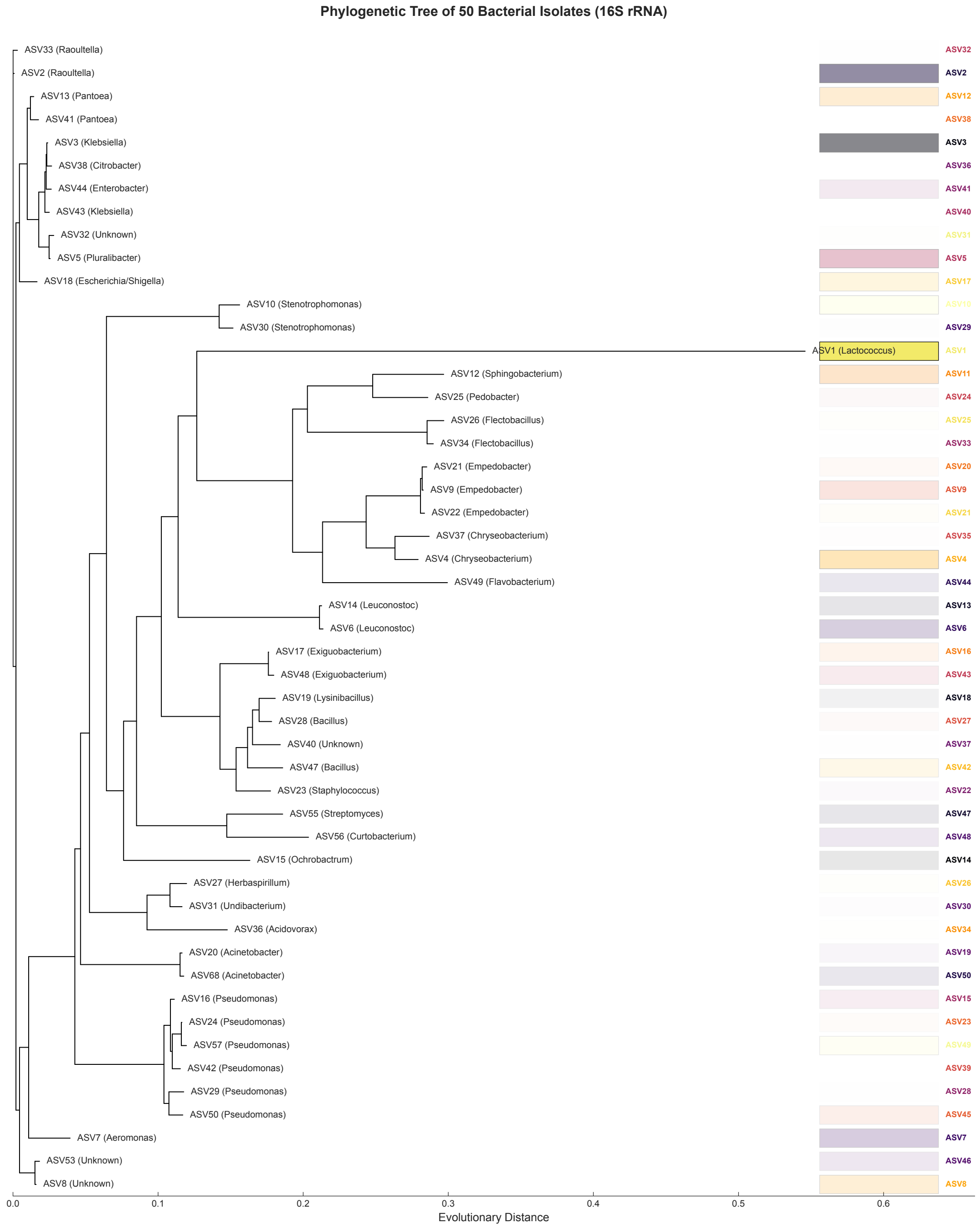


**Supplementary Fig. 1.** Phylogenetic tree of bacterial isolates. Maximum-likelihood phylogenetic tree based on full-length 16S rRNA gene sequences of the 54 bacterial isolates used in this study. Tree was constructed using EMBL-EBI tools[1]. The isolates span three phyla (Proteobacteria, Firmicutes, Bacteroidota) and 29 families, representing broad phylogenetic diversity.

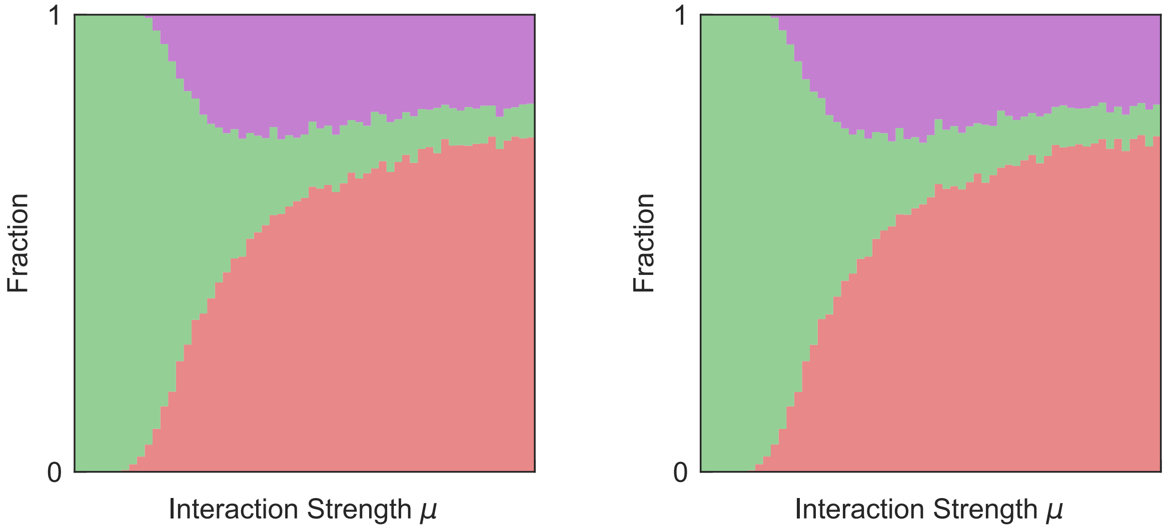


(a) Growth rate: mean 1, standard deviation 0.1

(b) Growth rate: mean 1, standard deviation 0.2

**Supplementary Fig. 2.** Phase diagrams with growth-rate heterogeneity. Coalescence outcomes when species have heterogeneous intrinsic growth rates sampled from normal distributions with mean 1 and standard deviations of **(a)** 0.1 and **(b)** 0.2. The qualitative transition from Mixture-dominated to Dominance/Restructuring outcomes with increasing interaction strength is robust to growth-rate variation.

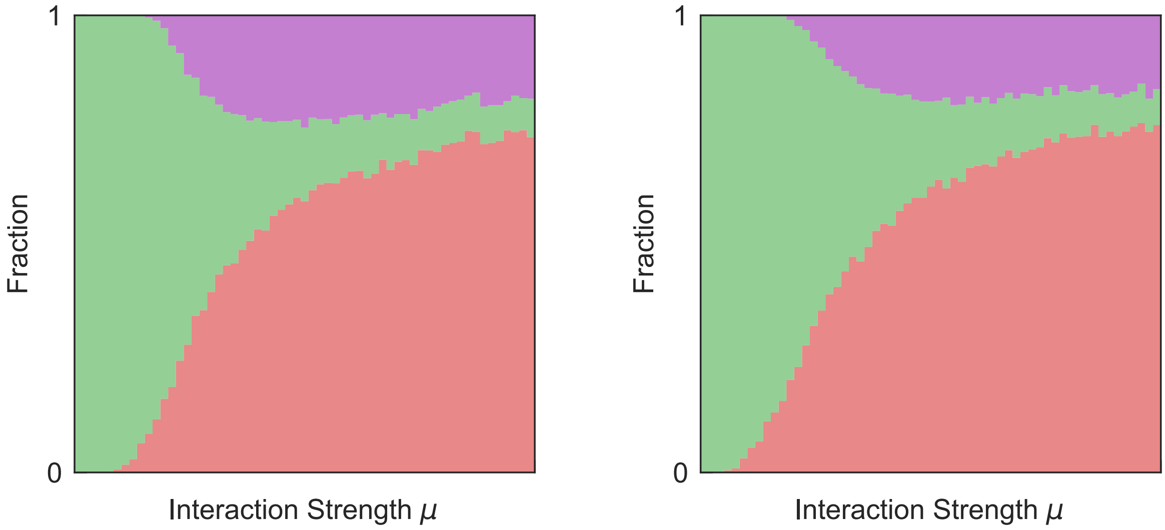


(a) Carrying capacity: mean 1, standard deviation 0.1

(b) Carrying capacity: mean 1, standard deviation 0.2

**Supplementary Fig. 3.** Phase diagrams with carrying-capacity variation. Coalescence outcomes when species have heterogeneous carrying capacities sampled from normal distributions with mean 1 and standard deviations of **(a)** 0.1 and **(b)** 0.2. The qualitative transition from Mixture-dominated to Dominance/Restructuring outcomes with increasing interaction strength is robust to carrying-capacity variation.

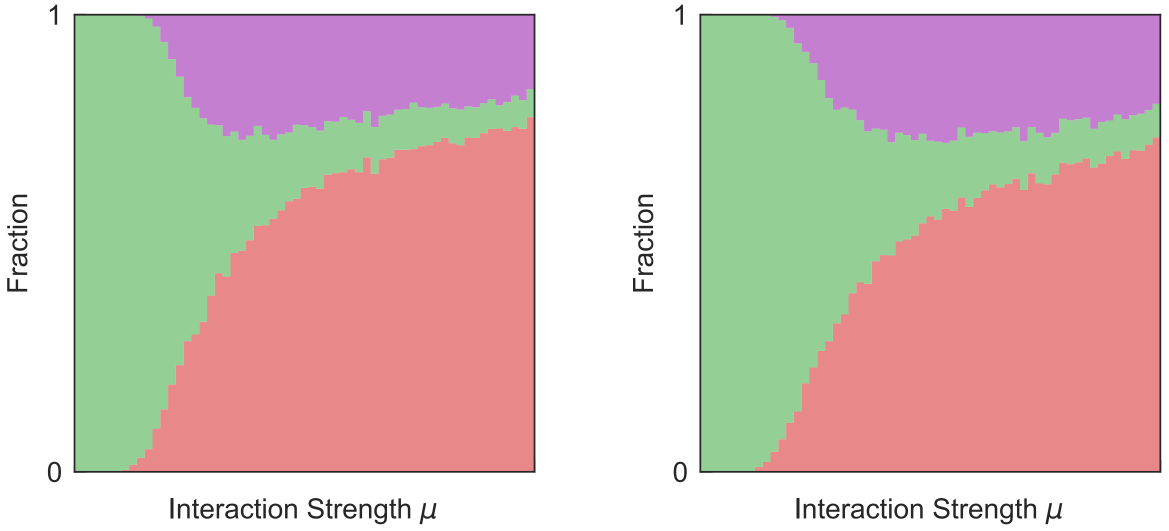


(a) Gaussian distribution (mean $\mu$, matched variance)

(b) Gamma distribution (mean $\mu$, matched variance)

**Supplementary Fig. 4.** Phase diagrams for alternative interaction coefficient distributions. Both distributions have the same mean and variance but different shapes. **(a)** Gaussian distribution (truncated at 0 to ensure non-negative coefficients). **(b)** Gamma distribution (inherently non-negative). The qualitative transition from Mixture-dominated to Dominance/Restructuring outcomes with increasing mean interaction strength $\mu$ is robust to the choice of distribution, confirming that the observed patterns depend primarily on interaction strength rather than distributional details.

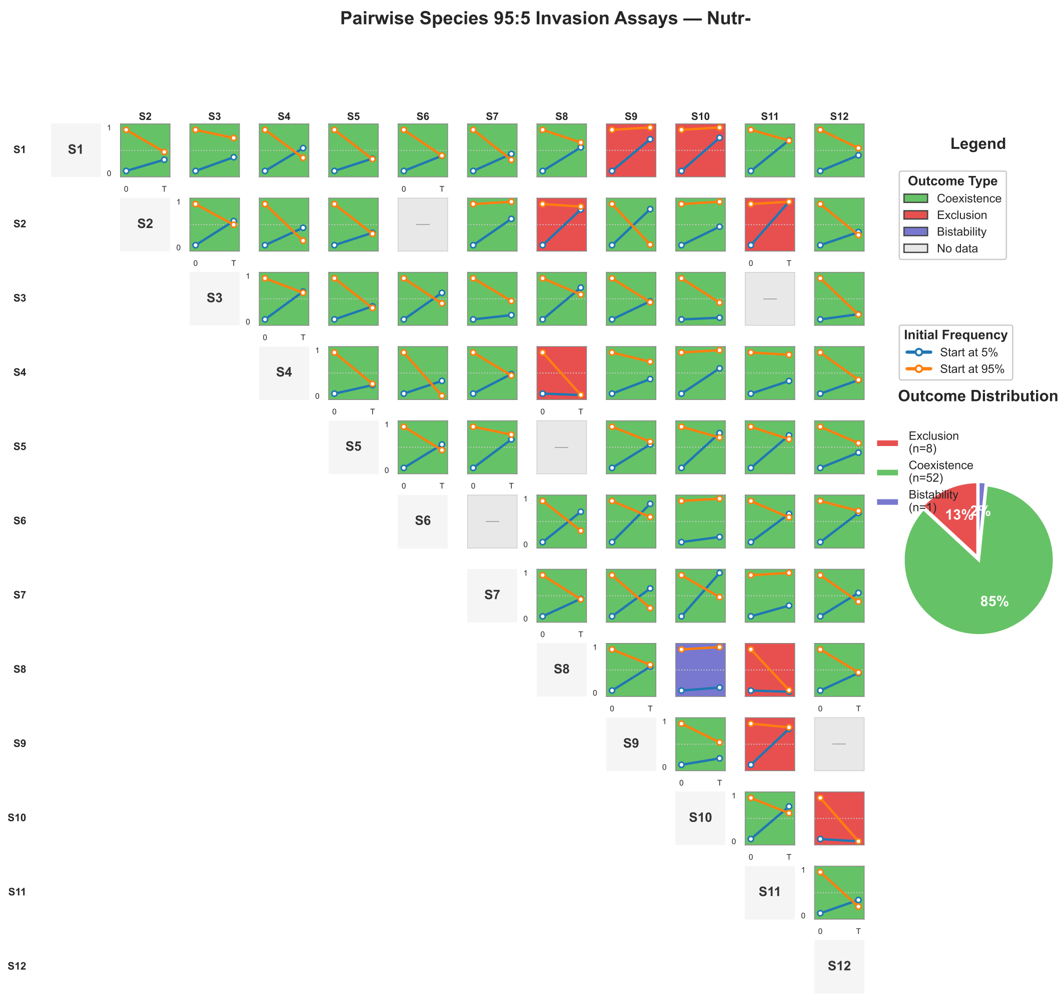


**Supplementary Fig. 5.** Pairwise invasion outcomes in Nutr− medium. Matrix showing competitive outcomes among the 12 most abundant isolates. Each cell indicates whether the invader (initially 5%) successfully established or was excluded after seven growth-dilution cycles. Low failed invasion frequency indicates weak competitive exclusion.

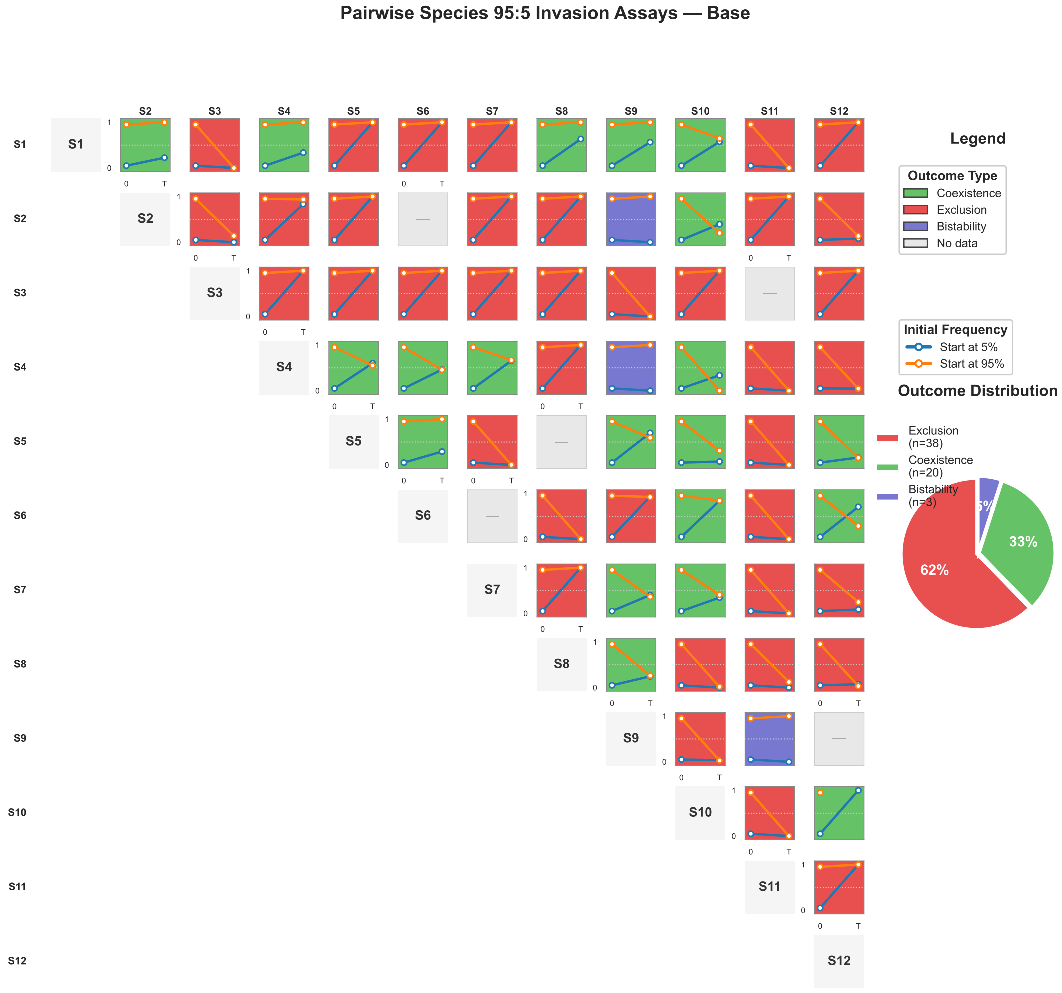


**Supplementary Fig. 6.** Pairwise invasion outcomes in Base medium. Matrix showing competitive outcomes among the 12 most abundant isolates. Each cell indicates whether the invader (initially 5%) successfully established or was excluded after seven growth-dilution cycles. Intermediate failed invasion frequency indicates moderate competitive exclusion.

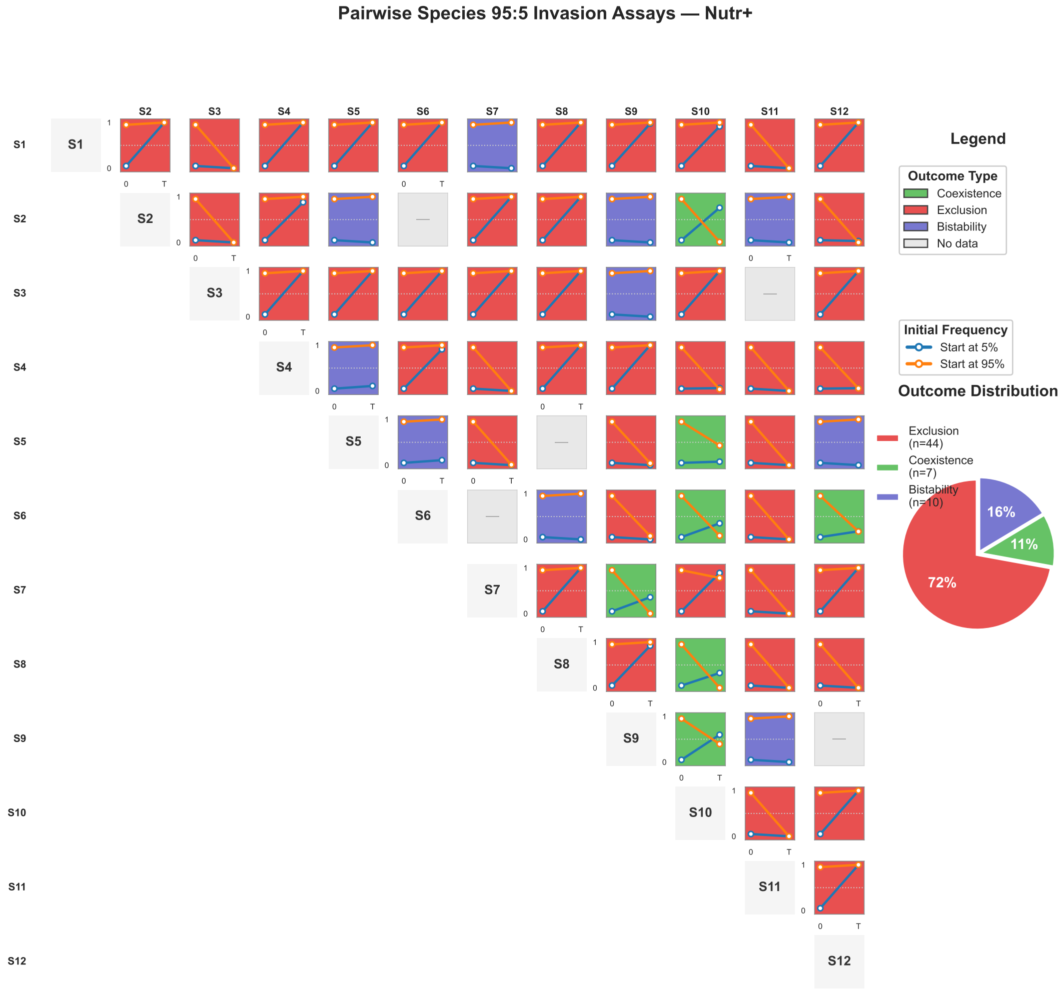


**Supplementary Fig. 7.** Pairwise invasion outcomes in Nutr+ medium. Matrix showing competitive outcomes among the 12 most abundant isolates. Each cell indicates whether the invader (initially 5%) successfully established or was excluded after seven growth-dilution cycles. High failed invasion frequency indicates strong competitive exclusion.

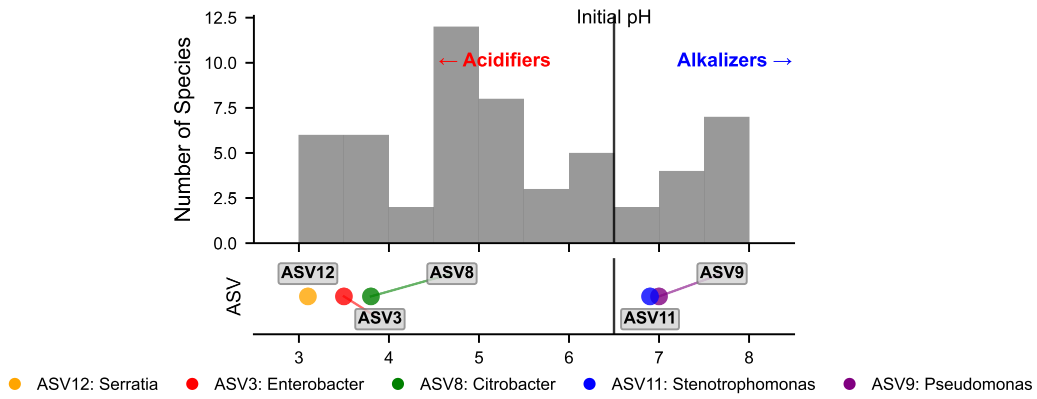


**Supplementary Fig. 8.** Monoculture pH modification by bacterial isolates. Distribution of monoculture pH after 15 h growth for all ASVs, showing the range of pH modification abilities relative to the initial pH of 6.5. Strong acidifiers (monoculture pH $< 5$) include ASV 12 (*Serratia*), ASV 3 (*Enterobacter*), and ASV 8 (*Citrobacter*). Species that maintain or slightly increase pH above initial (monoculture pH $> 6.5$) include ASV 11 (*Stenotrophomonas*) and ASV 9 (*Pseudomonas*). Bottom legend shows taxa identities for the labeled ASVs.

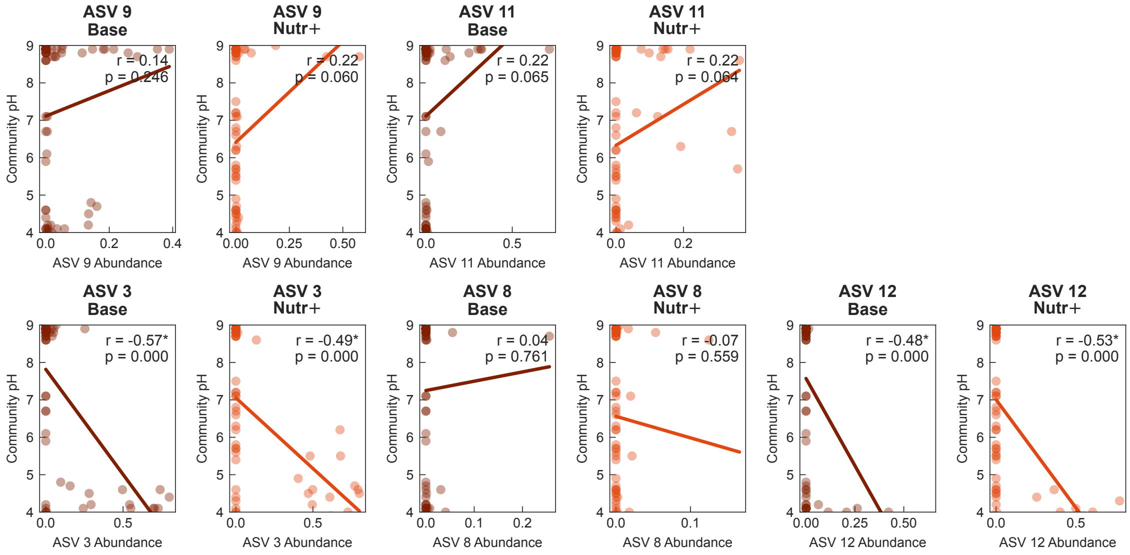


**Supplementary Fig. 9.** Dominant species class (acidifier vs. alkalizer) determines parental community pH. Scatter plots showing the relationship between dominant ASV relative abundance in parental communities and the resulting community pH at stabilization. Top row: alkalizers including ASV 9 (*Pseudomonas*) and ASV 11 (*Stenotrophomonas*), whose dominance is associated with higher community pH. Bottom row: acidifiers including ASV 3 (*Enterobacter*), ASV 8 (*Citrobacter*), and ASV 12 (*Serratia*), whose dominance is associated with lower community pH. For each ASV, left panel shows Base medium and right panel shows Nutr+ medium. Regression lines and correlation coefficients shown; asterisks indicate $p < 0.05$.

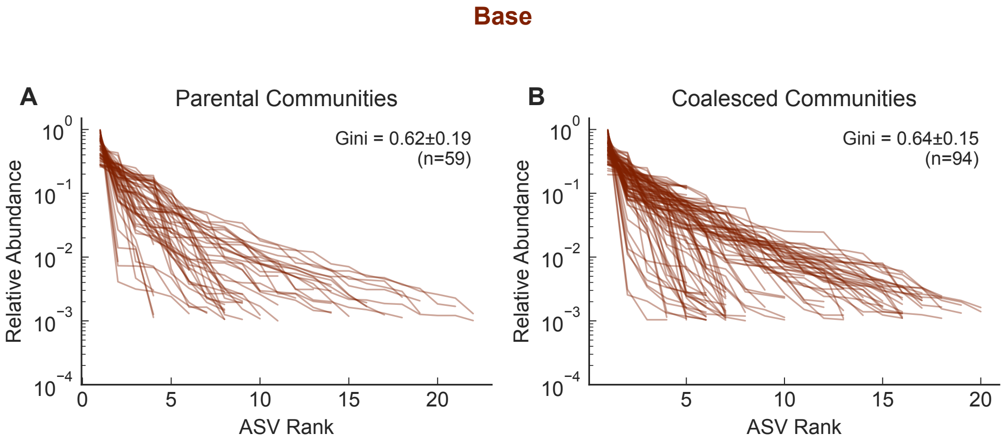


**Supplementary Fig. 10.** Rank-abundance curves comparing parental and coalesced communities in Base medium. **(a)** Parental communities before coalescence. **(b)** Coalesced communities after coalescence. Each line represents one community's rank-abundance curve. Gini coefficients quantify abundance inequality (0 = perfectly even, 1 = one species dominates). Parental: Gini = 0.62 ± 0.19 (n=59); Coalesced: Gini = 0.64 ± 0.15 (n=94). Both parental and coalesced communities exhibit substantial abundance skewness with similar Gini values.

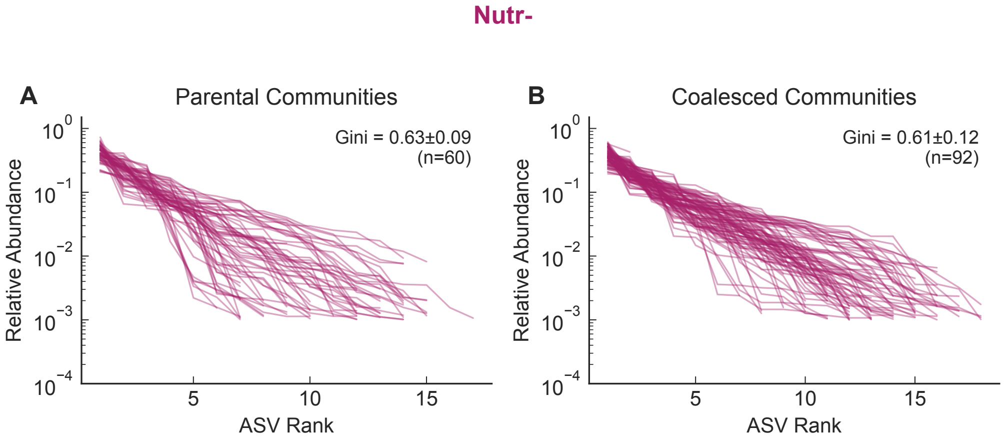


**Supplementary Fig. 11.** Rank-abundance curves comparing parental and coalesced communities in Nutr− medium. **(a)** Parental communities before coalescence. **(b)** Coalesced communities after coalescence. Each line represents one community's rank-abundance curve. Parental: Gini = 0.63 ± 0.09 (n=60); Coalesced: Gini = 0.61 ± 0.12 (n=92).

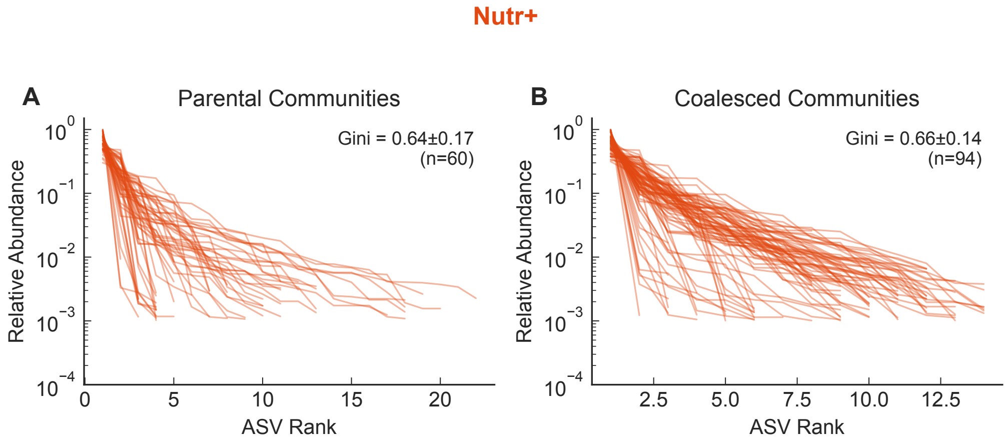


**Supplementary Fig. 12.** Rank-abundance curves comparing parental and coalesced communities in Nutr+ medium. **(a)** Parental communities before coalescence. **(b)** Coalesced communities after coalescence. Each line represents one community's rank-abundance curve. Parental: Gini = 0.64 ± 0.17 (n=60); Coalesced: Gini = 0.66 ± 0.14 (n=94).

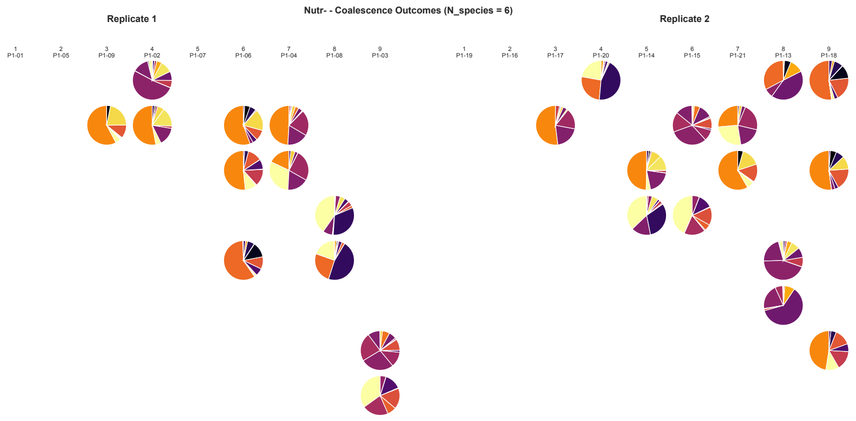


(a) Initial richness: 6 species

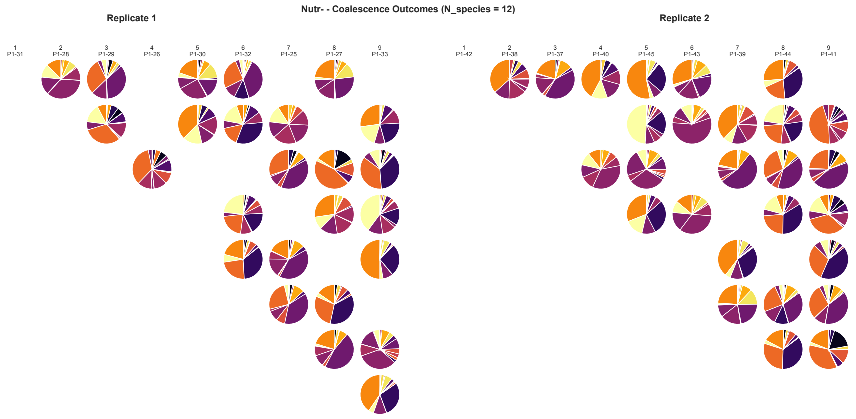


(b) Initial richness: 12 species

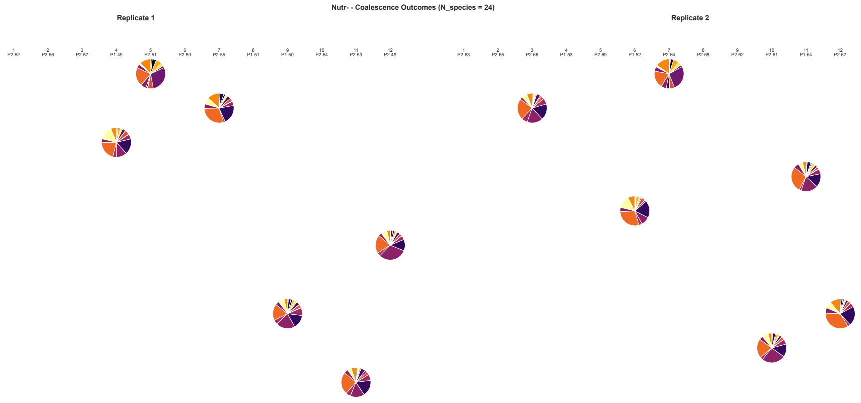


(c) Initial richness: 24 species

**Supplementary Fig. 13.** Coalescence outcome matrices in Nutr− medium. Each matrix shows pairwise coalescence outcomes for all parental community combinations at different initial richness levels. Pie charts display the species composition of coalesced communities; colors match parental origins. Under weak interactions, outcomes show more balanced mixing.

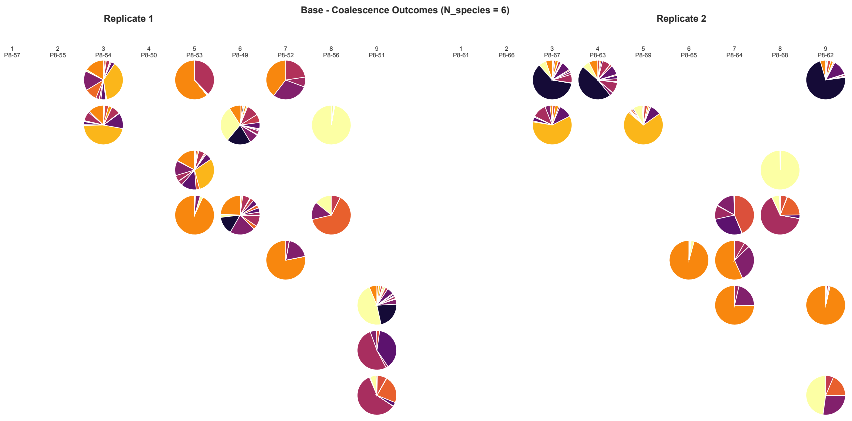


(a) Initial richness: 6 species

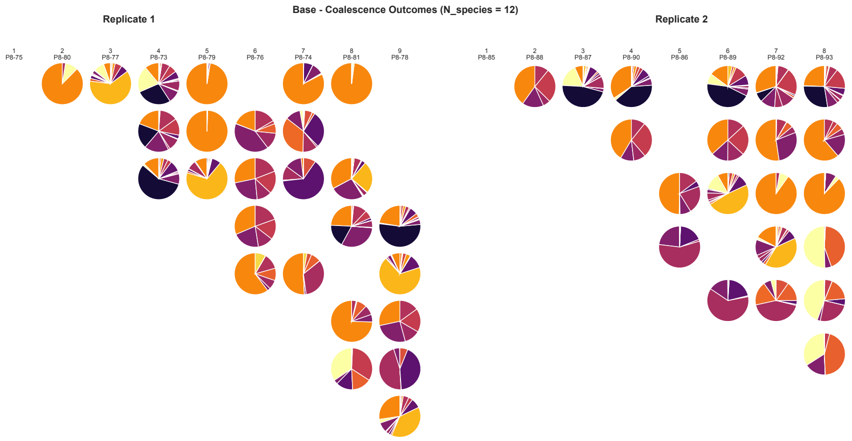


(b) Initial richness: 12 species

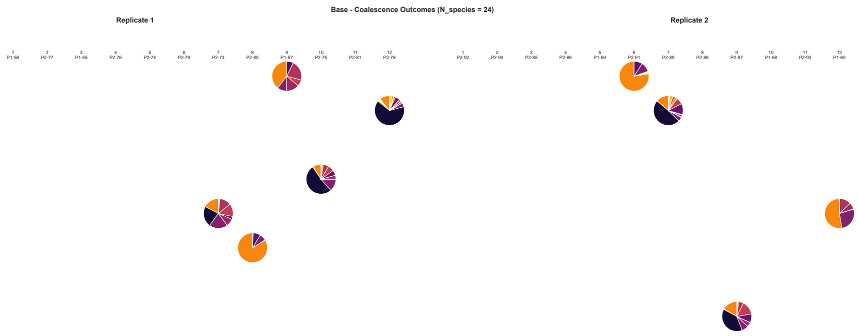


(c) Initial richness: 24 species

**Supplementary Fig. 14.** Coalescence outcome matrices in Base medium. Each matrix shows pairwise coalescence outcomes for all parental community combinations at different initial richness levels. Pie charts display the species composition of coalesced communities; colors match parental origins. Moderate interactions produce frequent one-sided outcomes (Dominance).

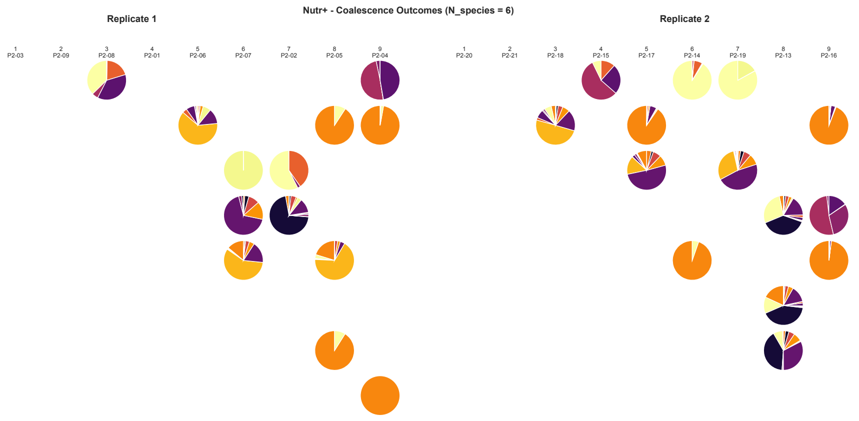


(a) Initial richness: 6 species

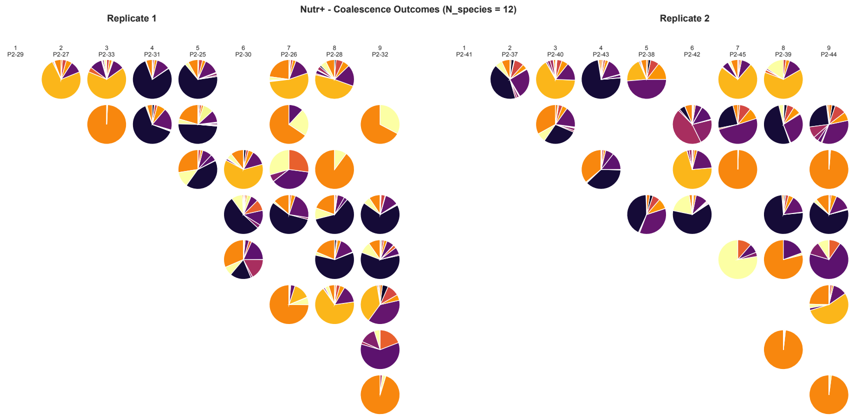


(b) Initial richness: 12 species

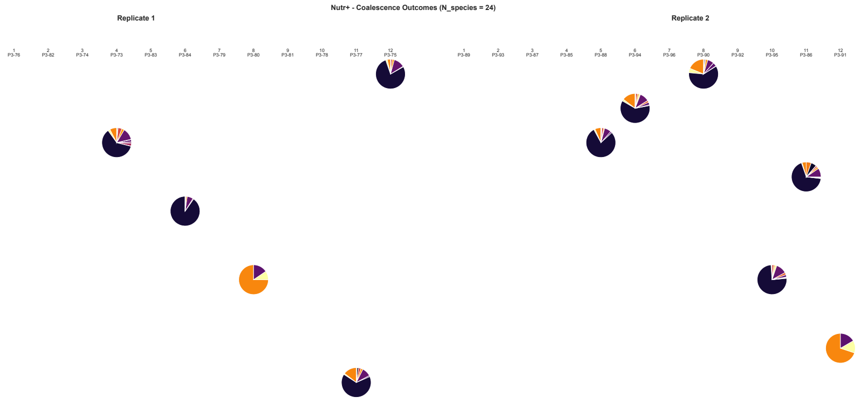


(c) Initial richness: 24 species

**Supplementary Fig. 15.** Coalescence outcome matrices in Nutr+ medium. Each matrix shows pairwise coalescence outcomes for all parental community combinations at different initial richness levels. Pie charts display the species composition of coalesced communities; colors match parental origins. Strong interactions produce predominantly one-sided outcomes (Dominance).

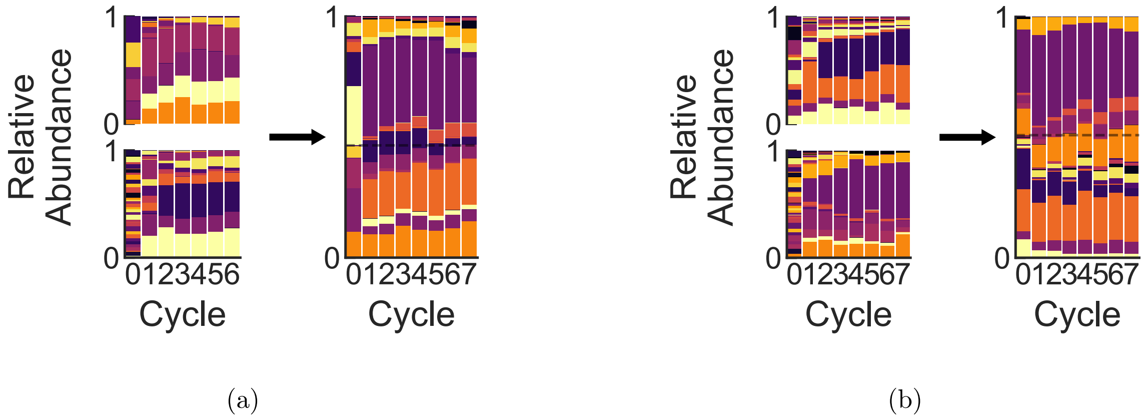


**Supplementary Fig. 16.** Time series of community composition during coalescence in Nutr− medium. Each panel **(a, b)** shows two parental communities (left, stacked vertically) and their coalesced outcome (right) over serial dilution cycles. Stacked bar charts represent relative species abundances at each time point. X-axis: serial dilution cycles (0–7); Y-axis: relative abundance.

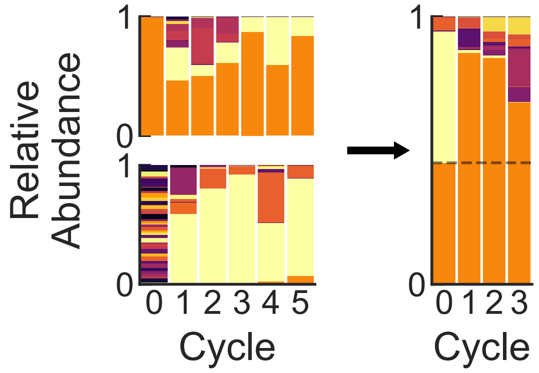


**Supplementary Fig. 17.** Time series of community composition during coalescence in Nutr+ medium. The panel shows two parental communities (left, stacked vertically) and their coalesced outcome (right) over serial dilution cycles. Stacked bar charts represent relative species abundances at each time point. X-axis: serial dilution cycles (0–7); Y-axis: relative abundance.

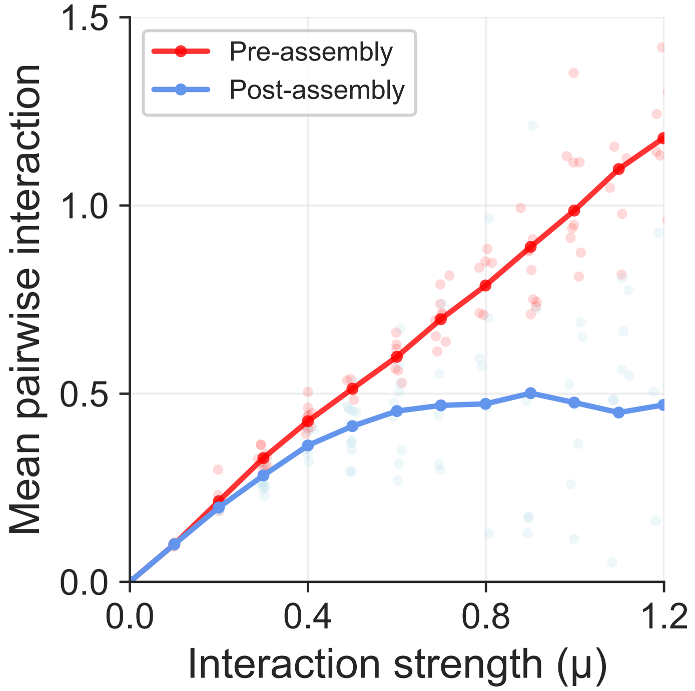


**Supplementary Fig. 18.** Assembly reduces mean pairwise interaction strength. Mean interaction strength before assembly (red, pre-assembly species pool) versus after assembly (blue, within-community) across different interaction strength parameters $\mu$. Points show individual simulation runs; lines connect means. Post-assembly communities consistently show reduced interaction strength, indicating that assembly filters out strongly competing species.

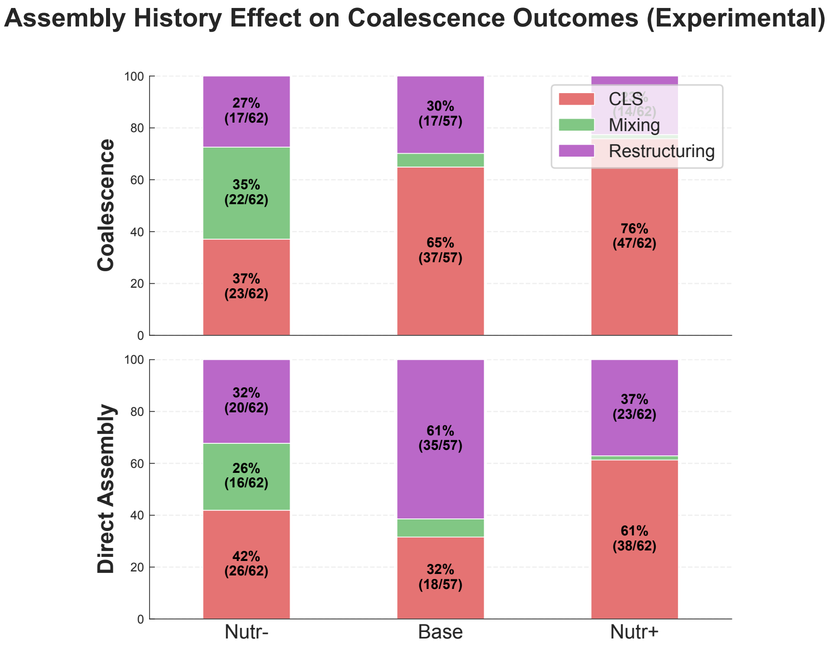


**Supplementary Fig. 19.** Assembly history promotes Dominance in experiments. Stacked bar plots comparing outcome distributions between coalescence (top row) and direct assembly (bottom row) across three nutrient conditions (Nutr−, Base, Nutr+). Percentages and counts (n/total) are shown for each category. In Base and Nutr+ conditions, coalescence shows elevated Dominance rates compared to direct assembly, consistent with simulation predictions (Extended Data Fig. 7, Supplementary Fig. 18).

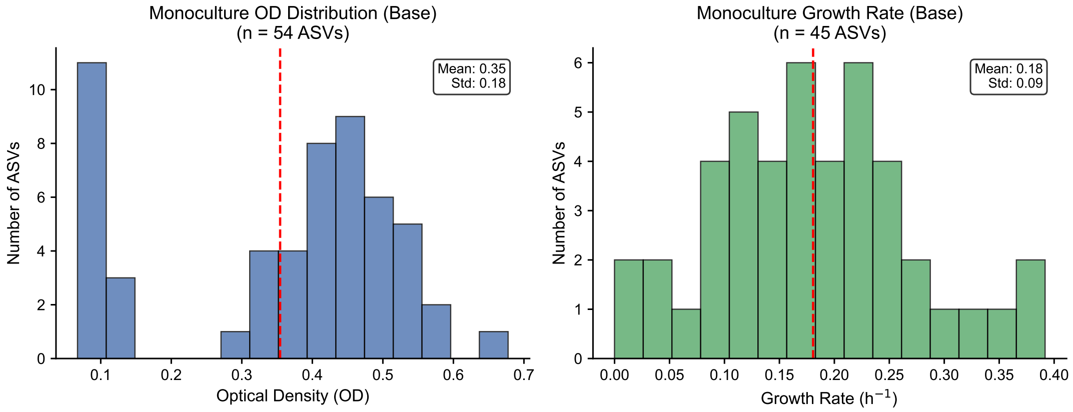


**Supplementary Fig. 20.** Phenotypic diversity of bacterial isolates in monoculture (Base medium). **(a)** Distribution of optical density ($OD_{600}$) reached by bacterial isolates during monoculture growth in Base medium ($n = 54$ isolates). **(b)** Distribution of exponential growth rates across isolates ($n = 45$ isolates with measurable growth rates out of 54 total). Growth rates were computed by log-linear regression of OD versus time during the exponential phase from full kinetic time-course measurements (475 cycles, ∼3 min intervals). The isolates exhibit broad variation in both growth yield and growth rate, reflecting phenotypic diversity that may contribute to varied coalescence outcomes in community mixing experiments.

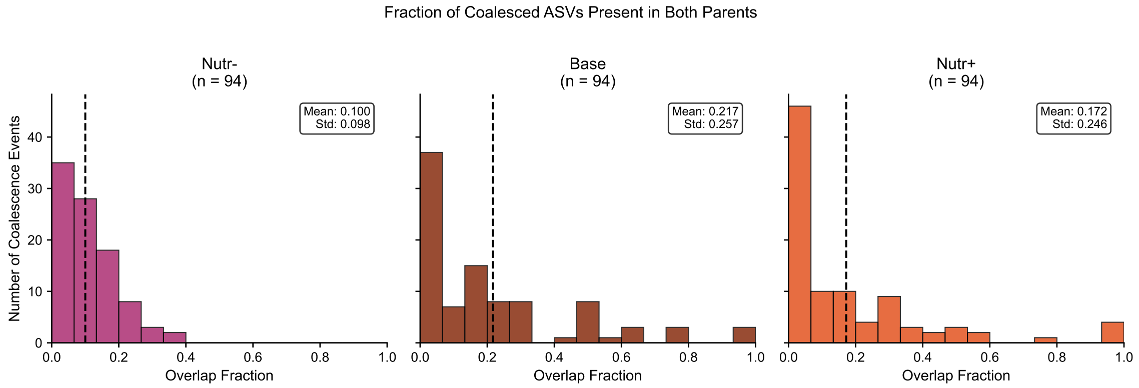


**Supplementary Fig. 21.** Fraction of coalesced ASVs present in both parental communities. Distribution of overlap fraction across coalescence events in three nutrient conditions. Overlap fraction is defined as the proportion of ASVs in the coalesced community that were present in both parental communities before mixing. Nutr−: mean = 0.10 ± 0.10 (n=90); Base: mean = 0.22 ± 0.26 (n=83); Nutr+: mean = 0.17 ± 0.25 (n=90). The relatively low overlap fractions across all conditions indicate that most surviving ASVs in coalesced communities originated from only one of the two parental communities, consistent with the prevalence of Dominance outcomes.

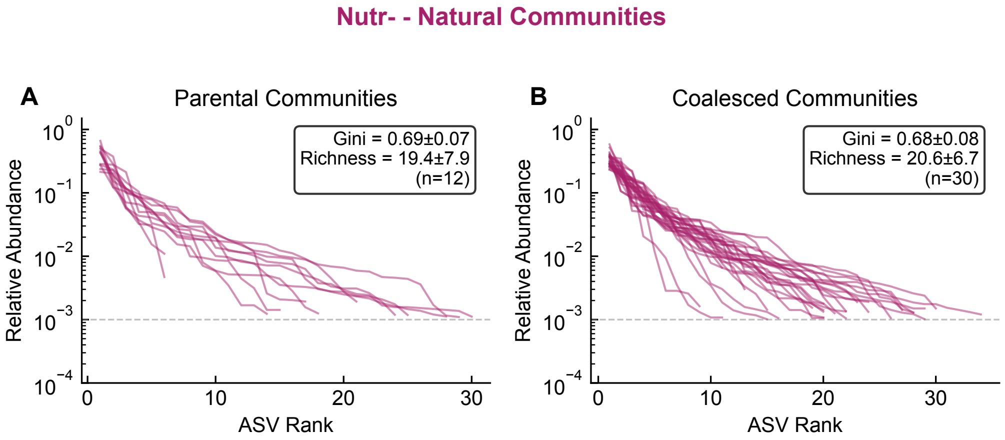


**Supplementary Fig. 22.** Rank-abundance curves for natural sample-derived communities in Nutr− medium. **(a)** Parental communities before coalescence. **(b)** Coalesced communities after coalescence. Each line represents one community's rank-abundance curve. Gini coefficients quantify abundance inequality (0 = perfectly even, 1 = one species dominates). ASV richness indicates number of ASVs above 0.1% relative abundance threshold. Horizontal dashed line indicates 0.1% threshold. Natural communities exhibit higher ASV richness compared to synthetic communities (which had controlled initial richness of 6, 12, or 24 species), reflecting the complex species assemblages in environmental samples.

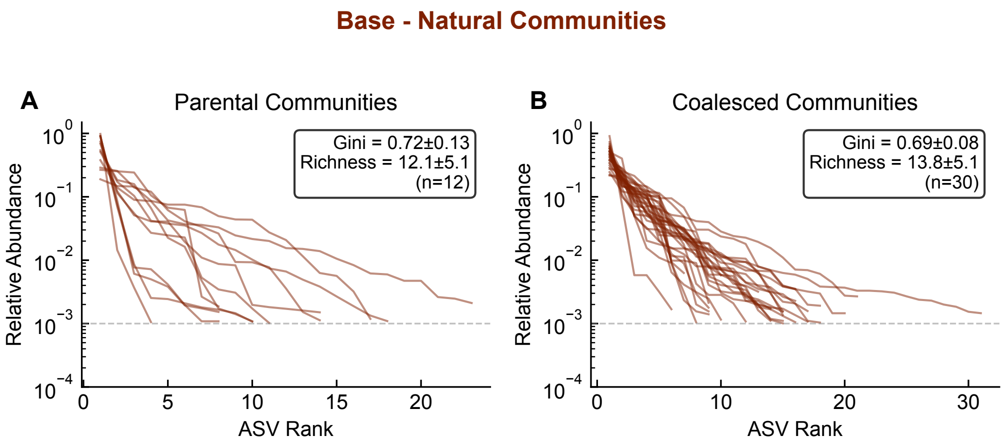


**Supplementary Fig. 23.** Rank-abundance curves for natural sample-derived communities in Base medium (natural-community analog of Supplementary Fig. 10 for synthetic communities). **(a)** Parental communities before coalescence. **(b)** Coalesced communities after coalescence. Each line represents one community's rank-abundance curve. Gini coefficients quantify abundance inequality (0 = perfectly even, 1 = one species dominates). ASV richness indicates number of ASVs above 0.1% relative abundance threshold. Horizontal dashed line indicates 0.1% threshold.

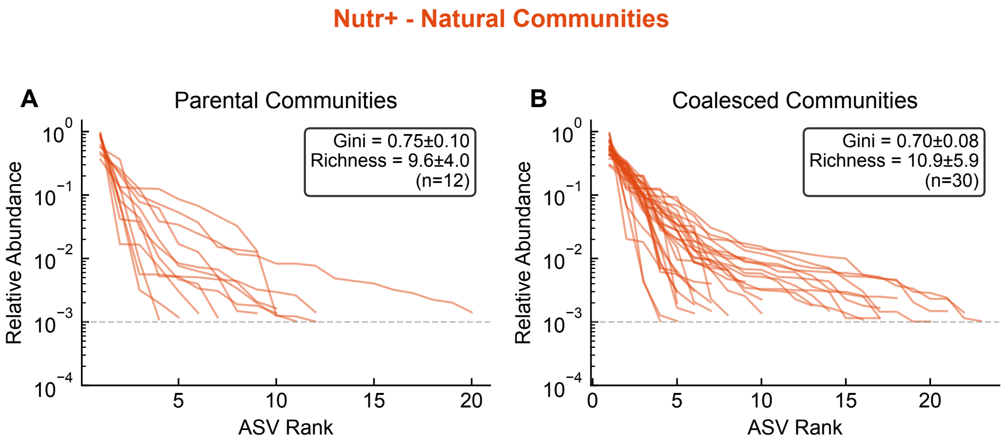


**Supplementary Fig. 24.** Rank-abundance curves for natural sample-derived communities in Nutr+ medium (natural-community analog of Supplementary Fig. 12 for synthetic communities). **(a)** Parental communities before coalescence. **(b)** Coalesced communities after coalescence. Each line represents one community's rank-abundance curve. Gini coefficients quantify abundance inequality (0 = perfectly even, 1 = one species dominates). ASV richness indicates number of ASVs above 0.1% relative abundance threshold. Horizontal dashed line indicates 0.1% threshold.

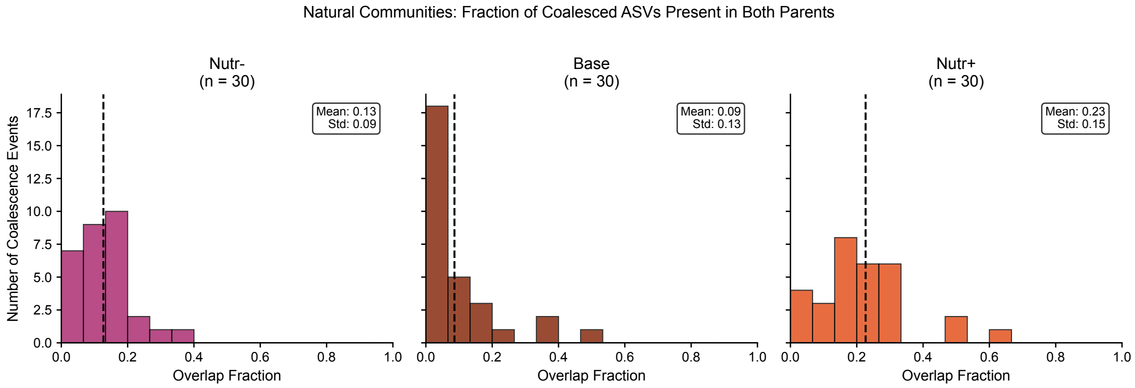


**Supplementary Fig. 25.** Fraction of coalesced ASVs present in both parental communities for natural sample-derived communities. Distribution of overlap fraction across coalescence events in three nutrient conditions. Overlap fraction is defined as the proportion of ASVs in the coalesced community that were present in both parental communities before mixing. Nutr−: mean = 0.13 ± 0.09 (n=30); Base: mean = 0.09 ± 0.13 (n=30); Nutr+: mean = 0.23 ± 0.15 (n=30). Similar to synthetic communities (Supplementary Fig. 21), the relatively low overlap fractions indicate that most surviving ASVs in coalesced communities originated from only one of the two parental communities, consistent with the prevalence of Dominance outcomes.

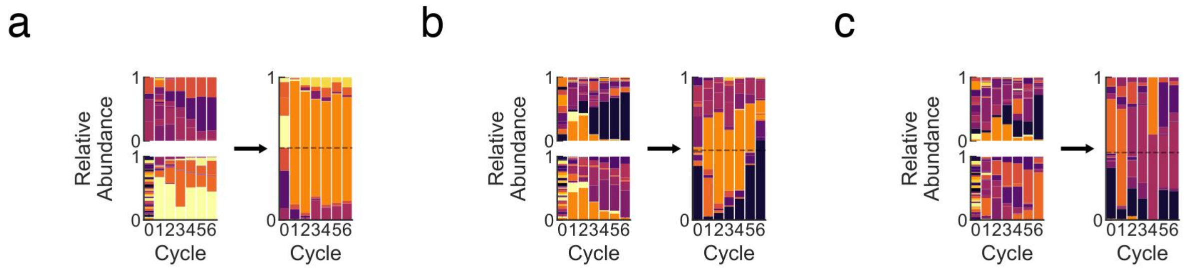


**Supplementary Fig. 26.** Time series of community composition during coalescence in Base medium. Each panel **(a–c)** shows two parental communities (left, stacked vertically) and their coalesced outcome (right) over serial dilution cycles. Stacked bar charts represent relative species abundances at each time point. X-axis: serial dilution cycles (0–7); Y-axis: relative abundance. These representative time courses illustrate the dynamics underlying Dominance and Restructuring outcomes, where one parental community rapidly displaces the other or the merged community converges toward a novel state.

### References

[1] Madeira, F. *et al.* Search and sequence analysis tools services from embl-ebi in 2022. *Nucleic Acids Research* **50**, W276–W279 (2022).
